# Sugarcane *bZIP16* regulates internode elongation by targeting gibberellic acid biosynthesis pathway

**DOI:** 10.64898/2026.10.12.758704

**Authors:** Saumya Shah, Sarvjeet Singh, Rashmi Gupta, Vaibhav Tewari, Shikha Jaiswal, Manoj Kumar Srivastava, Mehar Hasan Asif, Ajit Kumar Shasany, Manish Tiwari

**Author notes:** Academy of Scientific and Innovative Research, Ghaziabad, India (Sarvjeet Singh, Shikha Jaiswal, Mehar Hasan Asif, Manish Tiwari). Present Affiliation: Central University of Odisha, Koraput, India (Ajit Kumar Shasany).

## Abstract

Modern sugarcane cultivars are vegetatively propagated interspecific hybrids developed by extensive breeding and selection method to enhance sugar content, culm dimensions, desired plant architecture and disease resistance. Internode length greatly varies among existing sugarcane cultivars which determines culm length, biomass and sugar storage. Molecular factors responsible for this trait remains largely unknown in sugarcane. In this study, we characterized short and long internode sugarcane varieties and elucidated gene regulatory module interplaying with hormonal-network underneath the internodal growth. The genome size, pigment content and juice characteristics of short and long internode varieties differed with each other and corelated with gene expressions pattern. We established tissue and spatial transcriptome dataset, and comparative analyses demonstrated number of genes involved in cell cycle, cell expansion, transcription factors and hormone pathway including *bZIP16* are differentially expressed between short and long internode varieties. Biochemical, DNA-protein interactions and molecular evidences revealed that ScbZIP16 inversely related to GA content by repressing transcripts of *ent*-*isokaurene C2/C3-hydroxylase* and *GA20 Oxidase* in sugarcane. The further genetic evidences confirmed role of *ScbZIP16* in reducing GA biosynthesis pathway, flowering time and growth in *Arabidopsis*. This report of *ScbZIP16* regulatory module offers a potential candidate for sugarcane improvement programs.

**Summary Statement:** In this study, we found that Sugarcane *bZIP16* as a critical candidate for modulating GA biosynthesis which inversely regulates the internode length in sugarcane.

## 1 Introduction

Sugarcane, one of the most important industrial crops in the world, is mainly used for sugar and ethanol production. Sugarcane is also being utilized in sugar allied industries such as paper making, animal feed, organic manure and in chemical industry for the production of specialized compounds. Around 80 percent of global sugar production comes from sugarcane and 40 percent of fuel ethanol is being produced from the sugarcane (Li et al. 2025). Brazil is the highest sugarcane producing country accounts for 32.5 percent of the total global production in the year 2023-2024 followed by India (USDA, GAIN 2024). Sugarcane is cultivated in both tropical and sub-tropical regions of the globe including India (Dinesh Babu et al. 2022). Sugarcane, a representative of C4 plant, belongs to *Saccharum* genus which is a member of family Poaceae and tribe Andropogoneae. This genus comprises closely related species like *Saccharum barberi*, *Saccharum edule*, *Saccharum officinarum*, *Saccharum robustum*, *Saccharum sinense* and *Saccharum spontaneum* wherein interspecific crosses are possible, and this characteristic is commonly exploited by the breeders to develop new varieties (Mohan et al. 2022). The modern sugarcane varieties are interspecific hybrids of the domesticated sugar-producing species *S. officinarum* and wild relative *S. spontaneum* with multiple backcrosses to *S. officinarum* or to commercial-type hybrids. Genome of commercial sugarcane varieties is admixture of two distinct genomes having polyploidy and aneuploidy with estimated size is nearly 10 Gb (D’Hont et al. 1996). The first monoploid reference genome of R570 cultivar is about 382 Mb suggesting that polyploidization occurred in genome of both lineages, *S. officinarum* and *S. spontaneum* after their divergence (Garsmeur et al. 2018). Complete polyploid reference genome of the R570 cultivar showed distribution of 8.7 Gb sequences across 12 chromosomes (Healey et al. 2024). The allele-defined assembly of XTT22 cultivar and POJ2878 cultivar show extensive sub-genome recombination and non-homologous chromosomal rearrangements (Zhang et al. 2025; Wang et al. 2025). These genomic resources serve as foundation for accelerated improvement in sugarcane.

Breeding of domesticated cultivars with desired traits is a current method for sugarcane improvement, however, most of the traits such as high sucrose percentage, cane biomass and disease tolerance are polygenic and complex in nature regulated by multiple loci (Li et al. 2024). Further, sugarcane improvement breeding programs mainly focus on high sucrose level and increasing biomass. A recent GWAS study shows that sucrose content is a subject to strong artificial selection by breeders over the years and almost one-fifth of alleles in modern sugarcane cultivars are higher relative to previously grown varieties (Li et al. 2024). The genetic factors regulating mobilization of sucrose from source to maturing culm as well as redirection of brix to soluble sucrose was subject of extensive research in past years however it remains largely unknown till now. Several transcription factors associated with sucrose metabolism identified for their possible role in the processes comprising basic helix-loop-helix, homeobox knotted1-homeodomain genes, MYB and others in gene expression analysis (Papini-Terzi et al. 2009). As obvious, SWEET transporters seem to play very important role in long distance phloem transport of photosynthate and sucrose storage in parenchyma cells of internode (Mehdi et al. 2024). The sucrose content in cane increases at mature stage when vegetative growth slowed down, and lower internodes stored more sucrose compared to newly formed upper ones (Chandran and Premachandran, 2013).

Studies suggest that phytohormones such as auxin, gibberellin (GA), brassinosteroids and cytokinin plays an important role in tillering and internode growth by promoting cell elongation in sugarcane (Ren et al. 2023; Majda and Robert, 2018). The regulatory role of GA in promoting photosynthetic efficiency and plant growth is established in several studies (Wang et al 2025; Fan et al 2025; (Dayan et al. 2012). Earlier studies reported that foliar application of GA3 increases cane biomass, yield per hectare and sugar accumulation in sugarcane (Moore and Buren, 1978; Moore and Ginoza, 1980; Kuhnle et al. 1983;Roopendra et al. 2018). Recent transcriptome and gene expression studies demonstrate a differential expression of genes involved in sucrose metabolism, source–sink transport (Iqbal et al. 2011; Verma et al. 2017), tiller development and GA pathway observed upon GA3 application on sugarcane (Luo et al. 2023). Basic leucine zippers (bZIP) are major family of evolutionarily conserved transcription factors regulating pathogen defence, stress signalling, seed maturation and flower development in plant (Jakoby et al. 2002). The bZIPs play important roles in photomorphogenesis, cell elongation, energy metabolism, hormone, sugar signalling and reproductive development in *Arabidopsis* (Dietrich et al. 2011; Kang et al. 2010). Members of bZIP gene family like *bZIP16* integrates light and hormone signalling pathways to regulate early seedling development and altered expression of set of photosynthesis associated genes in *Arabidopsis* (Hsieh et al. 2012; Lindback et al. 2023).

Due to diverse uses, sugarcane demand is increasing globally and improvement in sugarcane yield through innovative approaches including genetic engineering is a pressing necessity. Stalk length is one of the key agronomic traits of varietal improvement programs and lot of variation occurs among sugarcane cultivars. A study shows that stalk length can reach up to 19.5 feet in height in some sugarcane variety (Du et al. 2026). The internodal elongation is a significant step of stalk growth that directly impacts cane height, sugar storage and overall biomass. Nevertheless, genetic and molecular mechanism responsible for culm height, and internode development is almost unknown in sugarcane. In this study, we thoroughly characterized short (CoLk14201) and long (CoLk16202) internode varieties, and genes expression changes in node and internode measured by comparing spatial transcriptome. A set of identified genes seem to be associated with distinct morphological characteristics of these varieties, and precisely expression of *ent*-*isokaurene C2/C3*-*hydroxylase* (*ScentIKH*) and *ScGA20 Oxidase* play a critical role in regulating internode length in sugarcane. We further revealed that ScbZIP16 negatively regulates GA biosynthetic genes expression leading to altered GA3 level in sugarcane varieties. The expression of *ScbZIP16* in *Arabidopsis* suppressed the transcripts of GA biosynthetic genes and active GA3 level resulted less seedling growth and delayed flowering time. Our findings establish *ScbZIP16* as a negative regulator of internode length and highlight its use in sugarcane improvement program in future.

## 2 Materials and Methods

### 2.1 Plant materials and sample collection

The two sugarcane varieties (CoLk14201 and CoLk16202) used in the study were grown at the farm of Indian Council of Agricultural Research-Indian Institute of Sugarcane Research (ICAR-IISR), Lucknow, India, located at 26° 56′N,80°52′E and 111 metre above sea level. Node and internode samples of both varieties have been collected at early (three months, stage 1; S1) and mature stage (nine-month, stage 2; S2). All the samples (nodes and internodes) were collected and immediately transferred into liquid nitrogen and kept in a −80 °C freezer. The sampling has been done quickly to minimize potential effects of fluctuations in the experiments.

The wild-type *Arabidopsis thaliana* ecotype Columbia-0 (Col-0) was used in this study hereafter referred as *Arabidopsis*. The Col-0 and transgenic lines were grown under standard conditions that is 16 hrs white light (115 µmol m^−2^ s^−1^) and 8 hrs dark at 22^0^C unless indicated otherwise. For dark and blue light induced hypocotyl growth assay, seeds were surface sterilized and cold-treated at 4 ^0^C for 2-3 days in the dark before germination. The seedlings were then grown in the continuous dark or blue light (50 µmol m^−2^ s^−1^) under long-day conditions {16 hrs white light (115 µmol m^−2^ s^−1^)} and 8 hrs dark photoperiod for the indicated days.

### 2.2 Photosynthetic pigments estimation and juice analysis

The photosynthetic pigments such as chlorophylls and carotenoids were extracted from 50 mg of leaf tissue in 5.0 ml DMSO. Total chlorophylls and carotenoids content were calculated using the following equations:

ChlA (μg/ g fresh weight) = 12.7 A663 −2.69 A645; ChlB (μg/ g fresh weight) = 22.9 A645 −4.68 A663, and Carotenoid (μg/ g fresh weight) = A480 +0.114A 663 - 0.638 A645. Analysis of sucrose percentage, brix, and purity coefficient in CoLk14201 and CoLk16202 have been carried out using the earlier described protocol (Rai et al. 2017).

### 2.3 Genome size determination by flow cytometry

The nuclear DNA content was estimated using the method described earlier with little modifications (Nogueira et al. 2022). Briefly, 20-30 mg of newly emerged young leaves of CoLk14201, CoLk16202 and experimental reference standard, *Pisum sativum* L. ‘Ctirad’ (9.09 pg) were chopped into small pieces containing chilled 500 µL Marie nuclei extraction buffer (Marie and Brown, 1993). The nuclei were stained by adding 15 μL propidium iodide solution (1 mg/mL) containing RNase and additives (PVP, βme, and triton x-100). The seeds of the external reference standard were provided by Dr. Jaroslav Doležel from the Laboratory of Molecular Cytogenetics and Cytometry, Institute of Experimental Botany, Olomouc, Czech Republic. Three samples comprising CoLk14201 and CoLk16202 and the external reference standard were used for the estimation using flow-cytometer (Cyflow Cube 8, Sysmex, Japan). The Nuclear DNA content was calculated in picograms (pg) using the formula described earlier by Doležel et al. (2007).

### 2.4 Total RNA isolation and transcriptome sequencing

Total RNA was isolated using the protocol described previously in (Furtado, 2014). This method involves the combination of the TRIzol kit and RNeasy Plant mini kit, in which initial extraction was done with guanidium thiocyanate-phenol-chloroform followed by filtration and resuspension of the RNA solution with a Qiagen RNeasy Plant Mini Kit (Qiagen, USA). RNA quantity, quality, and integrity were initially analysed using a NanoDrop 8000 spectrophotometer (Thermo Fisher Scientific, Wilmington, DE, USA). Final quality control before submission for sequencing was completed with an Agilent Bioanalyser 2100 (used with an Agilent RNA 6000 Nano kit: Agilent Technologies, Santa Clara, CA, USA). Sequencing libraries were generated from 3 μg total RNA using NEBNext Ultra RNA Library Prep Kit (NEB, USA). A paired-end (2x150 bp) library was constructed using the Illumina NovaSeq 6000 platform. A FASTP on the parameter of <Q20 was used for pre-processing the reads to get high-quality reads.

#### 2.4.1 Transcript profiling and differential expression analysis

The quality of raw RNAseq reads was assessed using FastQC v0.11.9. Subsequently, Trimmomatic v0.39 was applied to remove adapter sequences and trim bases with a Phred score below 20 (Bolger et al. 2014). The sugarcane reference genome available on the Saccharum Genome Database (Np-X.v20210328.Chr.fasta.gz) was used for indexing by Bowtie2-2.2.4. Cleaned reads have been mapped on the reference genome using Bowtie2-2.2.4. The alignment files were generated in SAM format and successively converted to BAM format and sorted using SAMtools v1.15 (Li et al. 2009). Aligned reads were assembled into transcripts using StringTie v2.1.7 (Pertea et al. 2016). Differential expression analysis was conducted using RStudio v4.3.1 and the edgeR v3.40.2 package (Robinson et al. 2010). Raw read counts were normalized, and differentially expressed genes (DEGs) were identified using a significance threshold of p < 0.005. Genes were classified as Upregulated if log_2_ fold change (log_2_FC) was 2 and downregulated if it was ≤−2.

For the estimation of stage-specific transcriptional differences following pair-wise contrasts were carried out in both the stages: CoLk14201NOD_vs_CoLk16202NOD, CoLk14201IND_vs_CoLk14201NOD, CoLk14201IND_vs_CoLk16202IND, CoLk16202IND_vs_ CoLk16202NOD (Supporting Information dataset S1). Apart from this, transcriptional differences between the stage 1 and stage 2 with the following pair-wise contrasts were also analyzed: CoLk14201INDS1_vs_CoLk14201INDS1, CoLk14201NODS1_vs_ CoLk14201NODS2, CoLk16202INDS1_vs_ CoLk16202INDS2 & CoLk16202NODS1_vs_ CoLk16202NODS2 (Supporting Information dataset S1).

#### 2.4.2 Functional Annotation

DEGs were functionally annotated for Gene Ontology (GO) terms, and KEGG pathways. AgriGO v2.0 (Tian et al. 2017) was used for GO enrichment analysis. KEGG pathway mapping was performed using KASS (KEGG Automatic Annotation Server) and KEGG Mapper.

### 2.5 Quantitative RT-PCR analysis

Total RNA (~2μg) isolated from node and internodes of CoLk14201 and CoLk16202 at S1 & S2 stages have been used for the synthesis of single strand cDNA (cDNA synthesis Kit, Thermo Scientific, USA). These cDNAs were used as templates for the quantification of relative mRNA levels with SYBR green (Excel Taq TM 2X Q-PCR Master Mix (SYBR, ROX, SMOBIO Technology, Inc.) in a Fast Real-time PCR system (StepOnePlus Real-Time PCR Systems, Applied Biosystems, USA). Gene specific primers have been designed through qPCR Primer & Probe Design Tool (https://eurofinsgenomics.eu/en/ecom/tools/qpcr-assay-design/) (Supporting Information Table S1). The PCR reactions for each sample were run in triplicate, and the mean value of each sample was statistically analysed. The 25S rRNA was used as an endogenous control (Iskandar et al. 2004).

For qRT-PCR analysis in *Arabidopsis*, approximately 100 mg tissue of mature rosette of Col-0 and transgenic lines were harvested and immediately frozen in liquid nitrogen. Total RNA of Col-0 and transgenic lines were isolated using the Spectrum™ Plant Total RNA Kit (Sigma-Aldrich) according to the manufacturer’s protocol, including on-column DNase I digestion (Qiagen) for 30 min at room temperature. The single strand cDNA synthesis and quantification of transcripts have been performed as mentioned in sugarcane (cDNA synthesis Kit, Thermo Scientific, USA). The *elongation factor 1 alpha* was used as an endogenous control (Supporting Information Table S1).

### 2.6 Gene cloning and yeast one hybrid interaction assay

The amplified PCR products of CDS region of Sc*bZIP16* and the promoter sequence of *Scent-IKH* and *ScGA20 Oxidase* (2kb; upstream from start codon) from CoLk16202 using High-Fidelity DNA Polymerase (NEB, UK) were ligated into the TA cloning vector pMD^TM^20 (Takkara, USA) by following the manufacturer’s instructions. The ligation mixtures were incubated overnight at 16°C. White colonies (transformed) were picked with the help of a toothpick, streaked on the LA plates (containing X-gal, IPTG, and ampicillin), and then inoculated in 5 ml LB containing ampicillin (100 mg/L) for isolation of plasmid DNA. The plasmid DNA was subjected to restriction digestion with the corresponding restriction enzymes and then analyzed on 0.8% agarose gel. Positive clones having an insert of appropriate size were sequenced using vector-specific M13F and M13R primers on a genetic analyser (Applied Biosystems, USA). The sequences obtained were analyzed for homology search in the NCBI database (http://www.ncbi.nlm.nih.gov) using blastx and blastp algorithm.

Further, *Scent-IKH* and *ScGA20 Oxidase* promoters was cloned in the pABAi vector and integrated into Y1H Gold yeast strain as described in manufacturer’s protocol (Takara Japan). The transformants were selected on synthetic defined (SD)–URA plate containing 200 ng/mL aureobasidin (AbA). The coding sequence of *bZIP16* has been cloned in prey vector pGADT7 (Takara, Japan). The effector constructs have been transformed into the *ScGA20 Oxidase*-pABAi and *Scent-IKH*-pABAi positive clone, separately. The positive clones were initially selected on SD-Leu plate for each construct. The spot assay has been used to know the promoter-effector interactions on SD-Leu (+200 ng/mL AbA) containing plate at indicated serial dilutions. The interactions of P53 and LAM with pT7 have been taken as positive and negative control, respectively in yeast spot assay.

### 2.7 Luciferase reporter assay and dual luciferase assay

For luciferase reporter assay, promoter of *Scent-IKH* and *ScGA20 Oxidase* were directionally cloned into pGreenII 0800-LUC vector whereas effector, *ScbZIP16* have been cloned into the pGreenII 62-SK vector, separately using restriction digestion method. The details of used primers for cloning are mentioned in supporting information Table S1. The constructs have been transformed into the *Agrobacterium tumefaciens* strain GV3101. The combination of promoter and effector constructs were transiently expressed in *Nicotiana benthamiana* leaves along with pSoup-P19 by following the protocol (Cheng et al. 2023). To visualize luciferase activity, luciferin solution (1 mM firefly D-luciferin, sodium-salt, Duchefa, in DMSO) sprayed on leaf surface, and image was captured under Molecular imager (ChemiDoc^TM^ XRS+, BIORAD, U.S.). The images of luciferase activity depicted with colour scales; blue indicating low activity and red indicating high activity.

The dual luciferase assay has been performed as described earlier (Hellens et al. 2005). Briefly, firefly luciferase (LUC) and Renilla luciferase (REN) reporter constructs have been agro-infiltrated in *N. benthamiana* leaves with pSoup-P19 as well. The leaf samples were harvested 48 hours post-infiltration and ground in liquid nitrogen. Around 100 mg sample was suspended in 100 µL luciferase extraction buffer (25 mM Tris-HCl, 2 mM EDTA, 10% Glycerol, 1% Triton X-100, 2 mM DTT, 1 mM phenylmethylsulfonyl fluoride; PMSF). The cellular debris were removed after 10 min centrifugation at 12,000 rpm (Thermo-Scientific sorvall ST 8R centrifuge), and supernatant was used in luciferase activity assay. The LUC and REN activities were analysed using Dual-Luciferase Reporter Assay System (Promega, Madison, WI, USA) according to the manufacturer’s protocol. The bioluminescence intensities were detected using the infinite M200 PRO TECAN (Switzerland).

### 2.8 Subcellular localization in *N. benthamiana*

For subcellular localization, *ScbZIP16* was cloned into the pENTR-D-Topo vector using a pENTR™ Directional TOPO^®^ Cloning Kit (Invitrogen, USA). After sequencing, entry clone was recombined in the destination vector pKF7WG2.0 using an LR Clonase (Invitrogen, USA). The details of used primers for cloning are mentioned in supporting information Table S1. The construct was introduced into *A. tumefaciens* (GV3101 strain) and infiltrated in four-week-old *N. benthamiana* leaves. Confocal imaging was performed using Leica Stellaris 8 confocal microscope (Leica Microsystems, Germany) 48 hrs post infiltration. Fluorescence of GFP was measured under the excitation at 488 nm and emission between 505–530 nm.

### 2.9 Gibberellic acid quantification

For the quantification of GA3, around 100 mg frozen nodes, internodes of mature cane, and 21-day old Col-0 and transgenic lines were ground in liquid nitrogen. The powdered samples were transferred to a 1.5 ml micro centrifuge tube and mixed with 1 ml chilled extraction solvent (80% methanol in 5% formic acid). After being vortexed for 2 minutes, the homogenised samples were extracted for 1h at 4°C with shaking. After extraction, the mixture was centrifuged at 10000xg for 10 min at 4°C. The supernatant was filtered with a 0.22 μm PTFE filter (Merck Millipore, IRL). Samples (5 μl) were then analysed by Triple Quadrupole 64595LC/TQ (Agilent Technologies, USA). Acquisition of mass was performed in negative ion mode as [M−H]^−^. In the mobile phase, 0.1% formic acid in water and 0.1% LC-MS grade acetonitrile was used as solvent A and solvent B, respectively, having the column oven temperature 30°C. Isocratic elution was performed in 65:35(V/V). Chromatographic separation was achieved on zorbax eclipse plus C-18 of 5 micron (analytical 4.6 mm x100 mm). Total run time, flow rate and the maximum pressure was 10 min, 0.3 ml min^−1^ and 1300 bar, respectively. The injection volume was 5 µl for authentic markers and samples. MassHunter acquisition software version 12.0 (Agilent Technologies, Santa Clara, CA, USA) was used to carry out all the operations, data collection, and data analysis. Gibberellic acid was measured in three different biological replicates. Quantification was done by the creation of calibration curves.

### 2.10 Generation of transgenic *Arabidopsis* lines

Complete CDS of *ScbZIP16* was amplified using forward and reverse gene-specific primers (GSPs, Supporting Information Table S1) containing *Xba*I and *Sac*I restriction sites, respectively. Amplicon was cloned between *Xba*I and *Sac*I sites into the pBI121 binary vector. Further, *ScbZIP16*-pBI121 overexpression construct was transformed into *Arabidopsis* (Col-0) background by floral dipping method (Clough & Bent 1998) using *A. tumefaciens* (GV3101). Transgenic plants were selected on ½ MS plate (0.22 g Murashige and Skoog media, 1% sucrose and 0.8% agar [pH 5.8]) containing 50 µg kanamycin /ml MS media. The independent lines exhibiting a 3:1 segregation ratio on kanamycin were selected for further propagation upto T4 generation and phenotypic analysis has been performed using homozygous lines.

### 2.11 Hypocotyl length measurement

For hypocotyl experiments, seeds of homozygous lines were surface sterilized using 70% ethanol and 2% Sodium hypochlorite containing Tween 20 (0.1 %) for 10 min and spread on half-strength MS (Himedia) supplemented with 1% sucrose, 0.5 mM 4-morpholineethanesulfonic acid (MES; pH 5.7), and 0.8% agar. The plates were stratified for three to four days at 4°C and then transferred in light (120 µmol m^−2^ s^−1^) for 16 hrs and 8 hrs dark photoperiod cycle. After the indicated times of growth and treatment, at least 25 to 30 seedlings were mounted on the agar plates, and digital pictures were taken using Nikon Camera (NIKKOR DX 18-140 mm/3.5-6.3f). Then, the hypocotyl lengths were measured using Image J software.

### 2.12 Protein extraction and immunoblot assay

For protein immunoblot assay, approximately 100 mg fresh tissues of *Arabidopsis* genotypes were harvested and ground to a fine powder using liquid nitrogen. Total protein extraction and western blotting was performed using the protocol described by Malakar et al. (2025) with some modifications. Total protein was isolated in 250 μL of extraction buffer [50 mM Tris-HCl (pH 8.0), 150 mM NaCl, 10% glycerol, 5 mM dithiothreitol (DTT), 1% NP-40, 0.5 mM phenylmethylsulfonyl fluoride (PMSF), and 1X ethylenediaminetetraacetic acid (EDTA)-free protease inhibitor cocktail] by centrifugation at 12,000 rpm for 10 min at 4°C. The protein concentration was estimated using Bradford assay. About 40 μg of total protein were denatured at 95°C for 5 min in 5X sodium dodecyl sulfate (SDS) sample buffer (100 mM Tris-HCl pH 7.5, 12% glycerol, 4% SDS, 200 mM DTT, 40 mM β-mercaptoethanol [ME], and 0.001% bromophenol blue), separated through one-dimensional SDS-PAGE (12% resolving gel, 4% stacking gel, 1.5 mm thick) and transferred to polyvinylidene fluoride (PVDF) membrane (Merck, Sigma) *via* wet transfer method. Further blots were blocked with 2% BSA in Tris-buffered saline plus Tween 20 for 1 h at RT, incubated overnight at 4°C with the respective primary antibodies, and subsequently incubated for 1 h at room temperature with horseradish peroxidase (HRP)-conjugated goat anti-rabbit (Invitrogen, 31460) secondary antibodies at a dilution of 1:10 000. Primary antibody for detecting |DELLA protein (Agrisera, AS111630, 1:1500) was used and chemiluminescence signals were detected using the Clarity MAXTM Western ECL substrate kit (BioRaD) through iBright^TM^ FL 1500 Imaging system (Invitrogen).

### 2.13 Statistical analysis

All the samples were collected in randomized manner, and experiments have been performed using biological and technical replicates as indicated in experiments and figure legends. Microsoft Office 2021, WA 98052-7329 USA (http://www.microsoft.com) was used for the calculation of mean values and standard differences. GraphPad prism software version 5.01, 225 Franklin Street. Fl. 26, Boston, MA 02110 (https://www.graphpad.com/) was used for the calculation of significant differences between the samples as *** (*p* < 0.001), ** (*p* < 0.01) and * (*p* < 0.05).

## 3 Results

### 3.1 Identification and characterization of short and long internode sugarcane varieties

In order to identify short and long internode varieties, around 20 popular sugarcane varieties, developed and released by ICAR-IISR, Lucknow, India for subtropical climatic conditions, have been examined for their internode length. For quantifying internode elongation, length of every internode from top to bottom of the mature cane has been recorded. Based on average length per cane, CoLk14201 and CoLk16202 have been identified as short and long internode producing sugarcane varieties and selected for further experiments (Figure 1). The data of average internode length suggested that CoLk16202 produces a significant larger internode compared to CoLk14201 (Figure 1C). The analysis of morphological characteristics such as cane height, girth, and number of nodes revealed a significant variation between CoLk14201 and CoLk16202 (Table 1). In terms of plant height, no significant difference was seen at early and mature stages between both the varieties (Figure S1 and Figure 1A, B; Table 1), however, number of nodes was significantly high in CoLk14201 (27 ± 4.03) compared to CoLk16202 (20 ± 2.13). In addition, cane girth, leaf fresh weight, root fresh weight and stalk fresh weight were significantly higher in CoLk14201 compared to CoLk16202 (Table 1). The morphological data showed that CoLk14201 has a greater number of nodes with significantly short internodes relative to CoLk16202.

**Figure 1.**
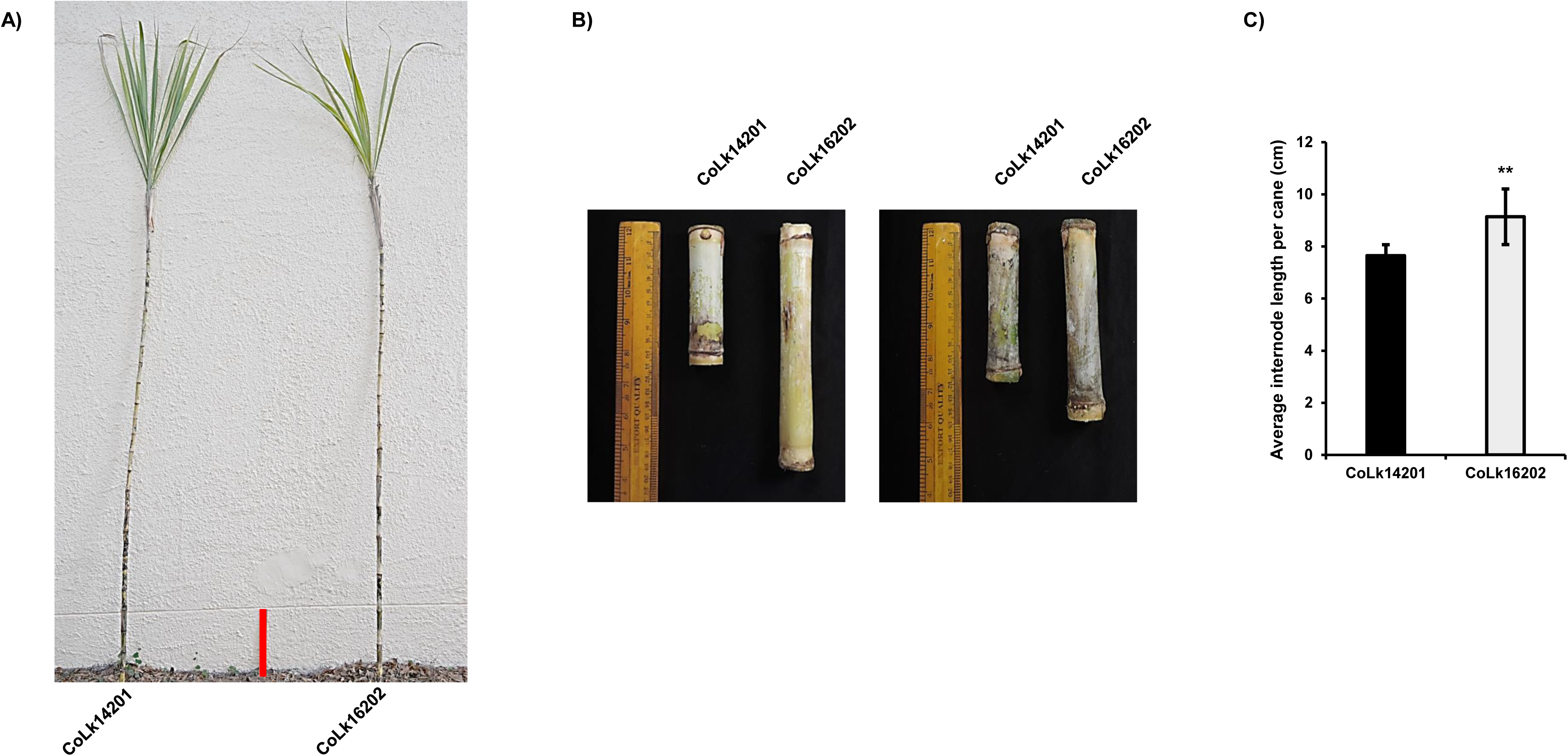
(A) Representative image of mature plant of sugarcane varieties displaying internode length variations (scale bar=30 cm in the middle). (B) Mature internode of sugarcane; length of fifth internode from top of the cane (left panel) and ground (right panel). (C) The average internode length per cane of sugarcane varieties. The bars represent data of seven individual cane. Asterisk (**) indicates *p*-value 0.001, level of significance calculated by paired student t-test.

**Table 1:**
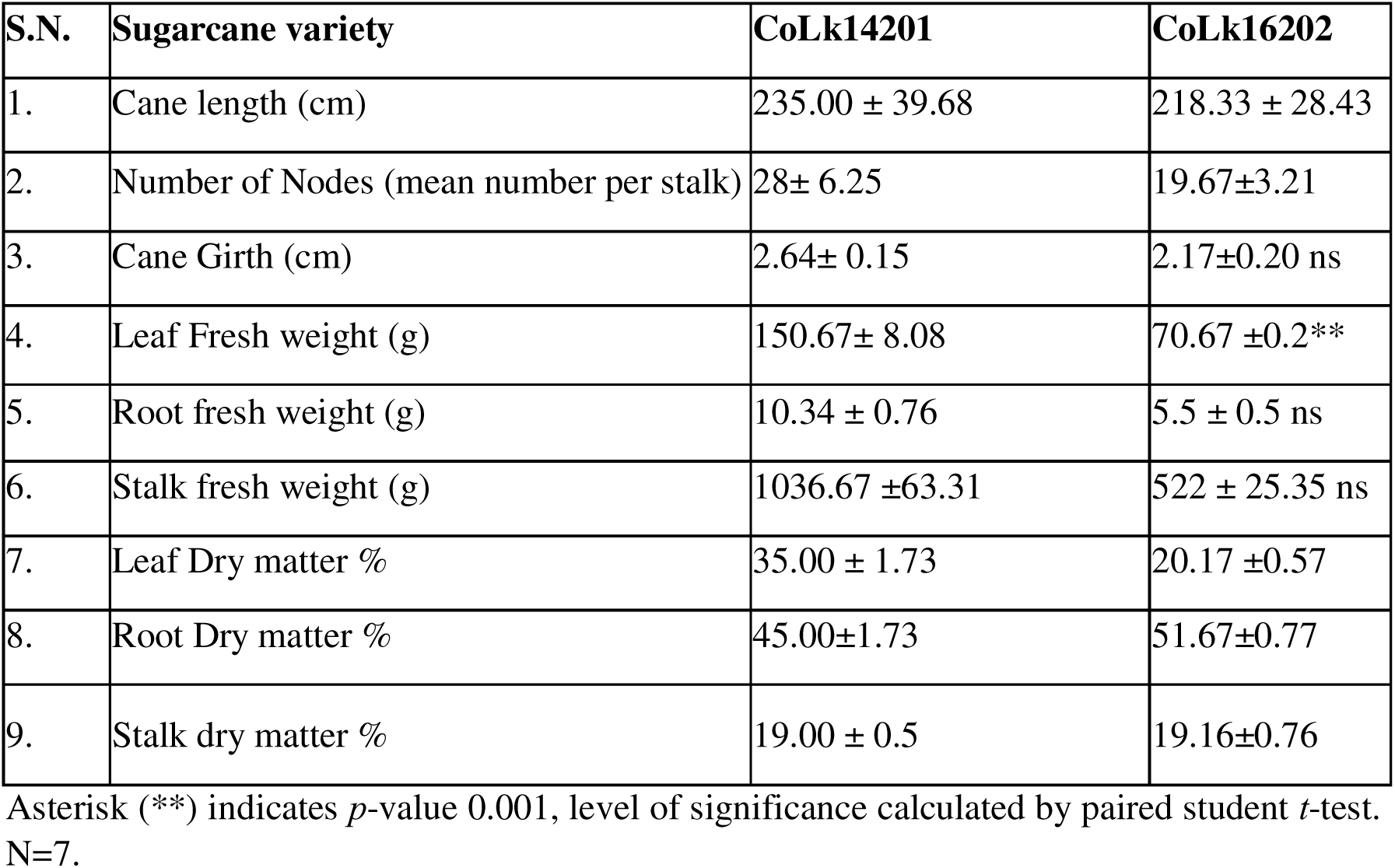
Morphological characteristics of mature plant of CoLk14201 and CoLk16202 varieties.

| S.N. | Sugarcane variety | CoLk14201 | CoLk16202 |
| --- | --- | --- | --- |
| 1. | Cane length (cm) | 235.00 ± 39.68 | 218.33 ± 28.43 |
| 2. | Number of Nodes (mean number per stalk) | 28± 6.25 | 19.67±3.21 |
| 3. | Cane Girth (cm) | 2.64± 0.15 | 2.17±0.20 ns |
| 4. | Leaf Fresh weight (g) | 150.67± 8.08 | 70.67 ±0.2** |
| 5. | Root fresh weight (g) | 10.34 ± 0.76 | 5.5 ± 0.5 ns |
| 6. | Stalk fresh weight (g) | 1036.67 ±63.31 | 522 ± 25.35 ns |
| 7. | Leaf Dry matter % | 35.00 ± 1.73 | 20.17 ±0.57 |
| 8. | Root Dry matter % | 45.00±1.73 | 51.67±0.77 |
| 9. | Stalk dry matter % | 19.00 ± 0.5 | 19.16±0.76 |

Asterisk (\*\*) indicates *p*-value 0.001, level of significance calculated by paired student *t*-test. N=7.

We determined genome size of CoLk14201 and CoLk16202 varieties through flow cytometry method. For this, newly formed leaves were used in flow cytometry and analysed the data. The results showed that nuclear DNA content of CoLk14201 and CoLk16202 was 10.54±0.165 pg and 10.34±0.035 corresponding to 10.3 Gb and 10.1 Gb, respectively (Dolezel et al. 2007). The obtained G1 peak of DNA in CoLk14201 and CoLk16202 with good resolution and coefficients of variation below 4.0 percent. The cells in G2 peak were prominent in reference but absent in sugarcane varieties indicating arrest of cell divisions (Figure 2A). We further examined photosynthetic activity of the sugarcane varieties by measuring photosynthetic pigments such as chlorophylls (ChlA and ChlB), and carotenoid in mature plant. The ChlA and ChlB content were found to be 229.76 ± 19.51, 54.08 ± 8.86 μg/g in CoLk14201 compared to 280.21±8.64 and 62.51±1.44 μg/g of fresh weight in CoLk16202 (Figure 2B). Similar trend was observed in case of carotenoid as CoLk14201 accumulated 15.34±1.11 μg/g relative to 19.81±0.74 80 μg/g of CoLk16202 suggesting a differential activity occurred in these varieties (Figure 2C). Moreover, we performed a detail juice analysis of varieties and data showed that CoLk16202 contained higher sucrose percentage as well as brix compared to CoLk14201 (Table 2). Together, CoLk14201 and CoLk16202 were morphologically distinct varieties and genome size of CoLk16202 was found to be ~200 Mb less than CoLk14201, which accumulated more photosynthetic pigments and sucrose.

**Figure 2.**
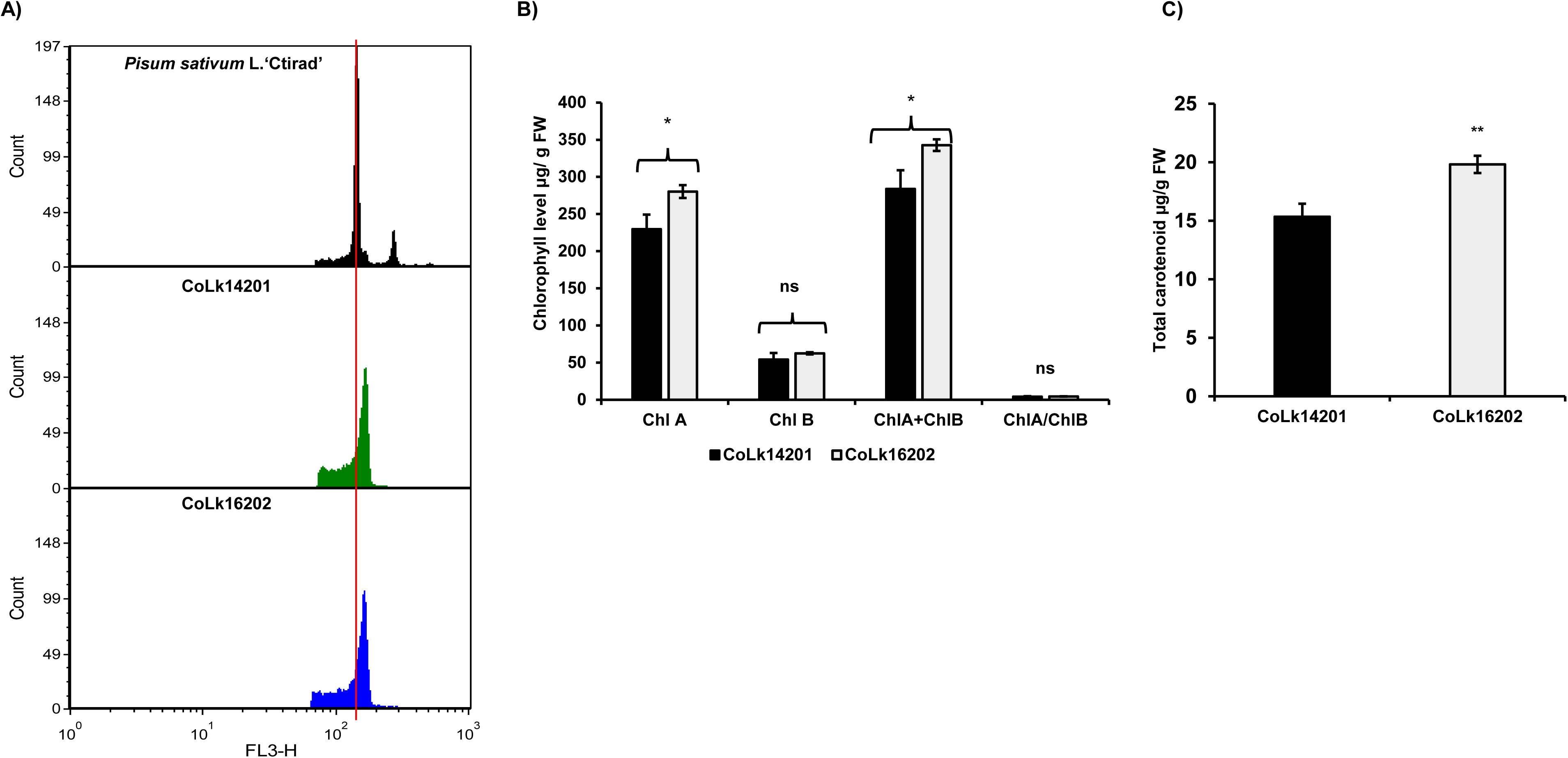
(A) Representative histograms of *P. sativum* L.‘Ctirad, CoLk14201 and CoLk16202. (B) Total chlorophyll content in µg/g fresh weight within CoLk14201 and CoLk16202. (C) Total carotenoid content (µg/g fresh weight) in CoLk14201and CoLk16202. The significant difference was compared from the mean ± SD of three independent biological replicates and three technical replicates. Statistical analysis was performed using one-way ANOVA (Dunnett’s multiple comparison test) for Total chlorophyll content analysis and paired student *t*-test for Total carotenoid content analysis. Significant difference: Asterisk (**) indicates *p* < 0.01, Asterisk (*) indicates *p* < 0.05, and ns indicates non-significant.

**Table 2:** Biochemical analysis of juice of sugarcane varieties.

| S.N. | Sample | Sucrose percentage | Purity coefficient | Brix |
| --- | --- | --- | --- | --- |
| 1. | CoLk14201 | 16.25±0.1979 | 85.331± 0.764 | 19.01±0.403 |
| 2. | CoLk16202 | 18.78±0.0141 ** | 85.33 ± 0.007 | 21.27± 0.0141 ** |

Asterisk (\*\*) indicates $p$ -value 0.001, level of significance calculated by paired student $t$ -test. N=7

### 3.2 Differential transcriptome analysis of node and internode of sugarcane varieties

To know the molecular factors responsible for internode elongation, comparative transcriptome analyses of node (NOD) and internode (IND) of CoLk14201 and CoLk16202 at two developmental stages i.e., early cane growth stage (S1; 3 months old) and mature cane (S2; 9 months old) of spring sown plants have been carried out. The RNAseq data revealed that 289 unigenes upregulated and 598 unigenes were downregulated at S1; and 1598 unigenes upregulated and 2808 unigenes were downregulated at S2 in the NOD of CoLk16202 (Figure 3; Table 3). Similarly, 629 unigenes upregulated and 149 unigenes were downregulated at S1 whereas 269 unigenes upregulated and 521 unigenes were downregulated at S2 in the IND of CoLk16202 (Figure 3; Table 3). The differentially expressed genes (DEGs) of same variety at S1 and S2 stages suggested that number of genes were found to be exclusively expressed in NOD and IND within varieties (Figure 3; Table 3). Among all the DEGs comparisons, we observed large number of unigenes (4148) downregulated at S1 in CoLk16202IND vs CoLk16202NOD, but only 412 unigenes downregulated at S2 (Figure 3). Among gene ontology (GO) function annotation of DEGs, we observed that *SWEET 6b* like bidirectional sugar transporters and *gibberellin-3 beta dioxygenases* were enriched in upregulated and downregulated comparisons, respectively of CoLk14201NOD vs CoLk16202 NOD at S1 (Figure S2). Similarly, several *SWEET 6b* like bidirectional sugar transporters were induced in the CoLk14201IND vs CoLk16202IND of S2 (Figure S2 & S3). It was seen that *gibberellin-3 beta dioxygenase* was found to be enriched in all the upregulated varietal comparisons (Figure S3). In addition, genes related to sugar metabolism, lignin synthesis, defence, stress related genes and large number of uncharacterized or unknown functions were also represented among all DEGs datasets comparison.

**Figure 3.**
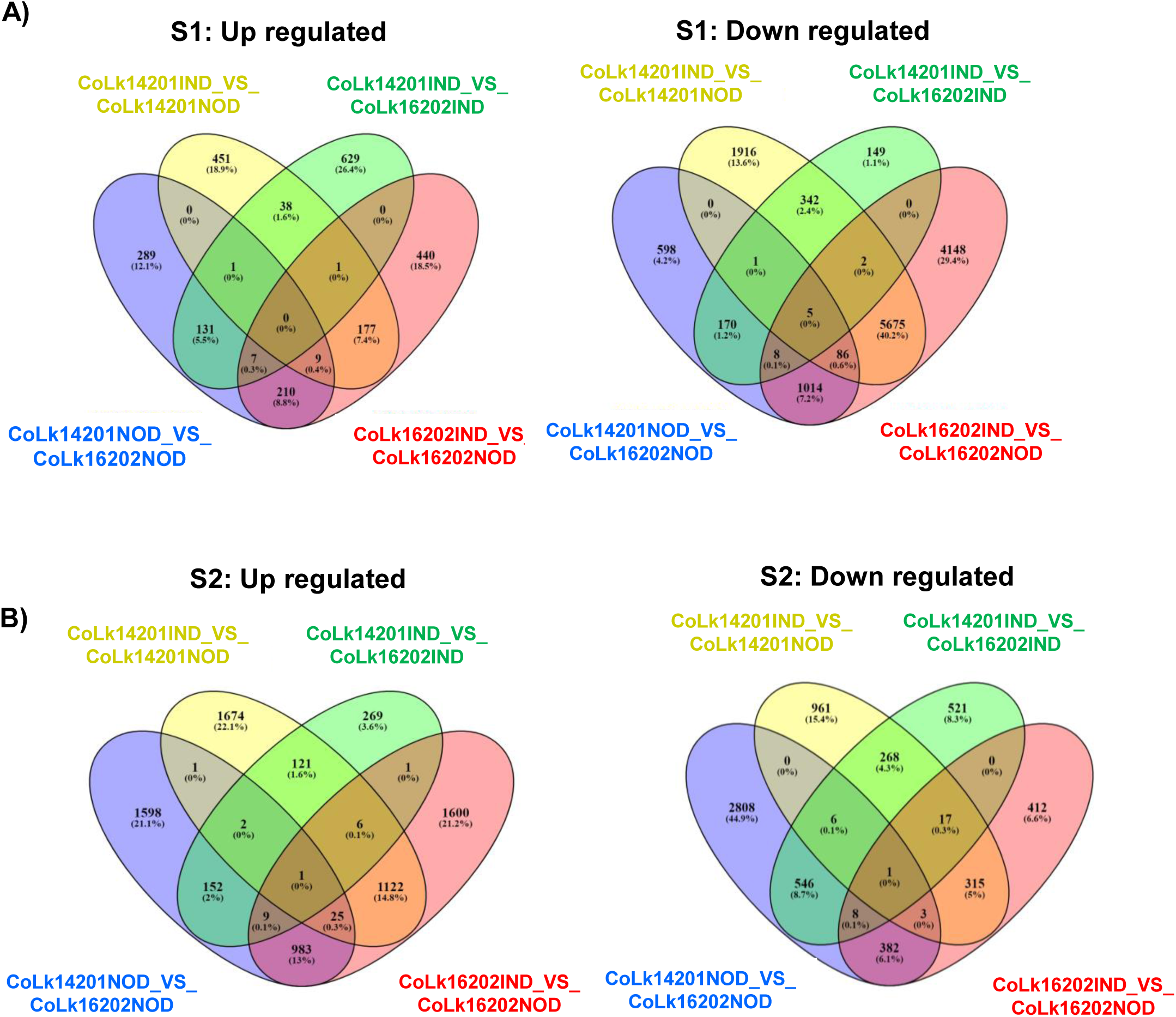
(A) Venn diagram common and unique differentially expressed genes in nodes and internodes of CoLk14201 and CoLk16202 at early stage (S1). (B) Venn diagram common and unique differentially expressed genes in nodes and internodes of CoLk14201 and CoLk16202 at mature stage (S2).

**Table 3:**
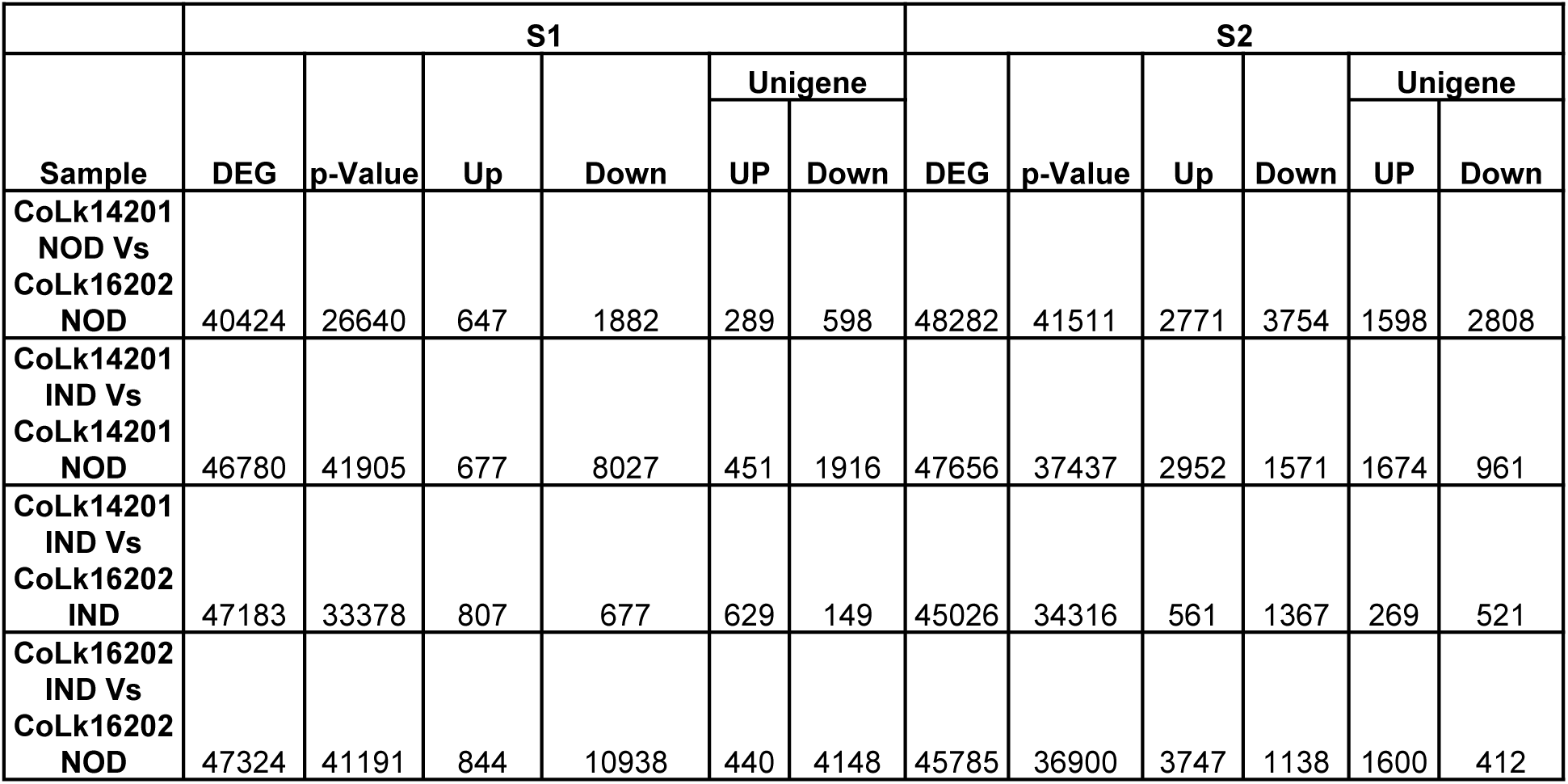
DEGs analysis in early (S1) and mature cane growth stage (S2) of CoLk14201 and CoLk16202.

| Sample | S1 |  |  |  |  |  | S2 |  |  |  |  |  |
| --- | --- | --- | --- | --- | --- | --- | --- | --- | --- | --- | --- | --- |
|  | DEG | p-Value | Up | Down | Unigene |  | DEG | p-Value | Up | Down | Unigene |  |
|  |  |  |  |  | UP | Down |  |  |  |  | UP | Down |
| CoLk14201 NOD Vs CoLk16202 NOD | 40424 | 26640 | 647 | 1882 | 289 | 598 | 48282 | 41511 | 2771 | 3754 | 1598 | 2808 |
| CoLk14201 IND Vs CoLk14201 NOD | 46780 | 41905 | 677 | 8027 | 451 | 1916 | 47656 | 37437 | 2952 | 1571 | 1674 | 961 |
| CoLk14201 IND Vs CoLk16202 IND | 47183 | 33378 | 807 | 677 | 629 | 149 | 45026 | 34316 | 561 | 1367 | 269 | 521 |
| CoLk16202 IND Vs CoLk16202 NOD | 47324 | 41191 | 844 | 10938 | 440 | 4148 | 45785 | 36900 | 3747 | 1138 | 1600 | 412 |

### 3.3 Genes related to cell division and growth differentially expressed in sugarcane varieties

The varietal transcriptome analyses indicated that numerous genes involved in cell division, expansion and cell wall loosening have been differentially expressed (Figure S4 & S5). Three *cyclin*-*dependent kinase A-2-like* genes (*CDKs*) were upregulated in IND of the CoLk16202. The CDKs, class of serine/threonine kinases together with cyclins, control G2/M phase transition and co-ordinating cell proliferation and differentiation (Guo et al. 2007). Numerous cyclins namely *cyclin-T1*-4-like, *cyclin-T1*-3-like and *cyclin-T1*-1 genes were significantly downregulated in CoLk16202 (Figure S4& S5). Reduced expression of *DRP5A* like genes has been noticed in NOD and as well as IND of CoLk16202 (Figure S5). Likewise, large number of *expansin*-like genes were also found to be downregulated in CoLK16202 in all the comparisons (Figure S4 & S5). Expansins promoted usually cell wall loosening, extension, cell enlargement, tissue softening, abscission and stress response in plant (Choi et al. 2006). Furthermore, several genes involved in lignin pathway such as *cinnamoyl-CoA reductase* (*CCR*) and *cinnamyl alcohol dehydrogenase* (*CAD*) represented in prevalent DEGs of NOD and IND (Figure S5D). The qRT-PCR analysis of selected genes such as *CCR, CAD* and *YABBY* followed the same expression pattern as seen in RNAseq experiment (Figure S5C and S5D; S8B).

The comparative transcriptomic data showed large number of transcription factors (TFs) differentially expressed among the most significant DEGs comprising *MYBs*, *WRKYs*, *bHLHs*, *basic leucine zipper* (*bZIP*), *B3-superfamily*, *NAC*, *AP2/ERF*, *FAR1*, *HOX*, *BEL1*, *CONSTANS* like 9, *BIM*, *TCP*, and *Cyclic DOF* (Figure S7, S8 & S9). The data showed that *MYBs*, *B3*, *Constans*, *bHLHs*, *ERFs*, *L10-interacting MYBs* (*LIMYBs*) and *Far1 like* were downregulated in NOD as well as in IND of the CoLK16202 variety (Figure S7). Among upregulated genes, *NACs*, *HOX 8 like* and *Cyclic DOFs* have been found to be common genes in both tissues. Upregulated TFs in the NOD of CoLk16202 at mature cane stage comprised *Erf118*, *bHLH13*, *MYBs* and *Constans* like indicating a significant varietal difference in genes expressions (Figure S8). Interestingly, higher expressions of various *DIVARICATAs*, *UNE12*, *ARF6-like*, *BIM2*, and *cyclic dof factor 2* have been observed in IND of CoLk16202 relative to CoLk14201 (Figure S9). Furthermore, growth-regulating factors (*GRFs*) regulate stem elongation, meristem maintenance, cell expansion and longevity in GA dependent manner (Omidbakhshfard et al. 2015). The higher expression of two *GRF*-like genes in IND of CoLk16202 corelating transcriptome data with morphological differences (Figure 4, Figure S9B). Together, gene expressions analyses showed a distinct tissue specific expression pattern of number of TFs involved in diverse regulatory processes occurred during early and mature cane development stages in studied sugarcane varieties.

**Figure 4.**
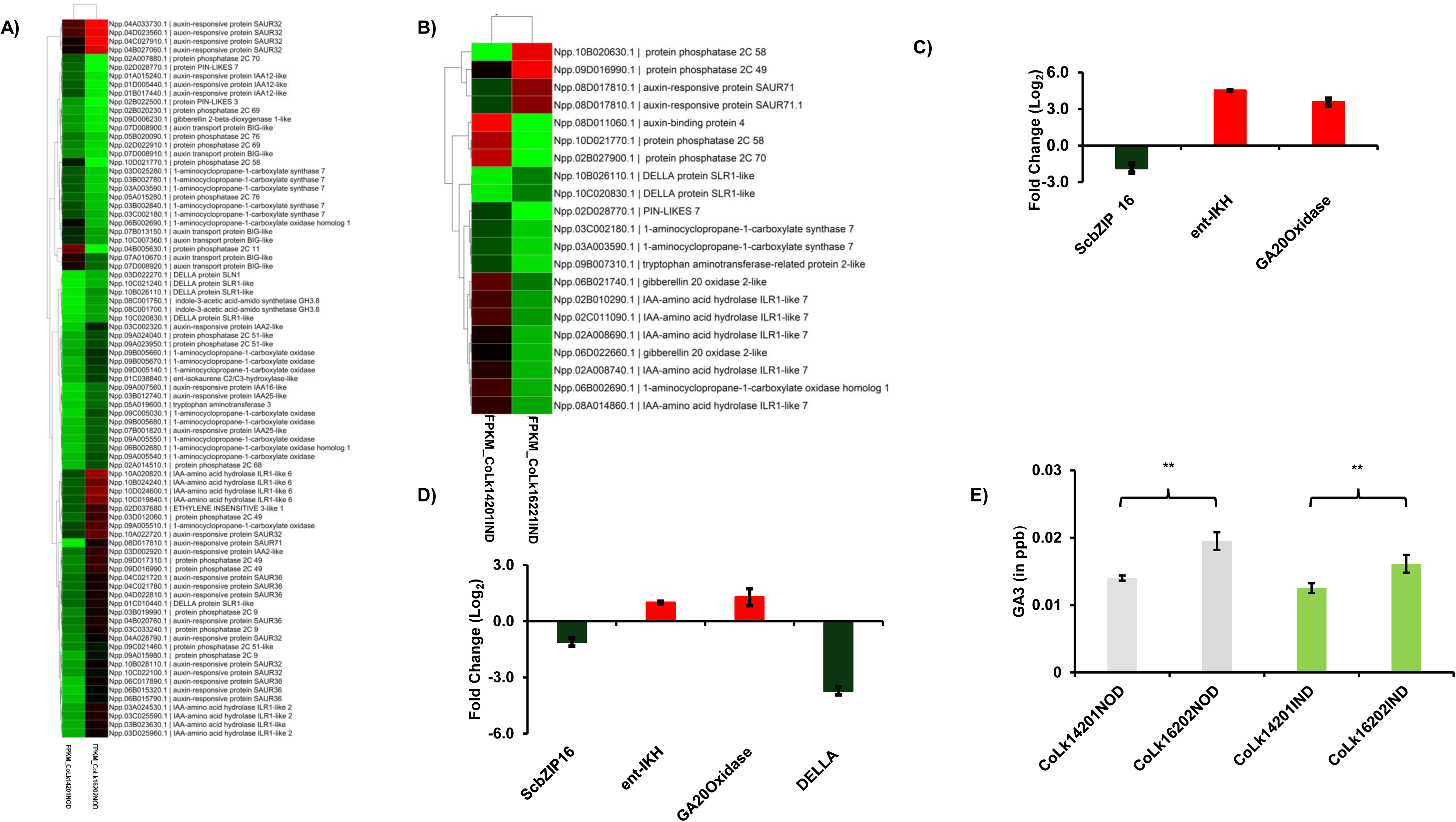
(A) Heatmap expression atlas of phytohormone related DEGs in the nodes of CoLk14201 and CoLk16202 at S2. (B) Heatmap expression atlas of phytohormone related DEGs in the internodes of CoLk14201 and CoLk16202 at S2. (C) Relative gene expression analysis of *ScbZIP16; ent*-isokaurene C2/C3-hydroxylase-like, *ent*-*IKH*; gibberellin 20 oxidase 2-like, *GA20 Oxidase*-like and *DELLA* related to GA biosynthesis in the node of CoLk14201 vs CoLk16202 which was analysed using pooled samples of three independent biological and three technical replicates (D) Relative gene expression analysis of *ScbZIP16* and *GA20 Oxidase*-like in the internode of CoLk14201 vs CoLk16202 comparison at S2 which was analysed using pooled samples of three independent biological and three technical replicates. (E) GA3 estimation (per 100mg) in node and internode of mature cane of CoLk14201 and CoLk16202 which was analysed using three independent biological and three technical replicates. The significant difference calculated using the one-way *ANOVA* (Bonferroni’s Multiple Comparison Test). Significant difference asterisk (*) indicated *p* < 0.05

### 3.4 *bZIP16* transcripts level inversely associated to internode elongation in sugarcane

Amongst significant DEGs, an interesting expression pattern of *bZIP16* has been observed in varietal comparisons of sugarcane. We noticed an increased expression of *bZIP16* in NOD and IND of the CoLk14201 found in RNAseq data relative to long internode carrying CoLk16202 (Figure S7 & S9). To confirm this, qRT-PCR analysis of NOD and IND of mature cane has been carried out and results validated previous observation by showing a significant reduction of *ScbZIP16* transcripts in both tissues of the CoLk16202 (Figure 4C and 4D). On the other hand, mRNA level of GA synthesis genes namely *Scent-IKH* and *ScGA20 Oxidase* were significantly higher in CoLk16202 compared to short internode sugarcane variety (Figure 4C&D). Furthermore, expression of *gibberellin 2 beta dioxygenases* and *DELLA SLR1* like transcripts have found to be altered in RNAseq as well as qRT-PCR data indicating a major contribution of GA pathway in cane elongation process (Figure 4). The *DELLA SLR1* like gene encode RGA protein, a central regulator of GA signalling, suggested a strong corelation between GA and internodal growth in sugarcane (Figure 4). We next sought to quantify the GA level in NOD and IND of sugarcane varieties. The LC-MS result showed a significant higher GA3 accumulation in the NOD and IND of CoLk16202 corresponding to CoLk14201 (Figure 4E). Together, expression profiling and GA3 measurement data suggested that bZIP16 can bind to promoters of GA pathway genes to regulate transcription of these genes in order to determine GA3 levels in sugarcane varieties.

### 3.5 ScbZIP16 localized in nucleus and suppressed transcription of GA biosynthesis genes

Inverse relation between *ScbZIP16* expression and GA pathway laid us to hypothesised that ScbZIP16 negatively regulates transcription of GA biosynthesis genes in sugarcane. Foremost, we determined subcellular localization of ScbZIP16:GFP tagged protein in mesophylls cells of *N. benthamiana*. The examination of GFP tracking clearly showed a nuclear localization of ScbZIP16 within leaf epidermal cells (Figure 5A). Further, TFs binding sites analysis within promoters of *Scent-IKH* and *ScGA20 Oxidase* cloned from CoLk16202 variety showed presence of multiple G-box motives including four core G-box motif (−241, −838, −1226, −1925 for *ScGA20 Oxidase*; and −1999, −1275, −804, −215 for *Scent-IKH*) where bZIPs can bind (Figure S10). To test ScbZIP16 binding on the promoter of *ScGA20 Oxidase* and *Scent-IKH*, we performed yeast one hybrid assay using ScbZIP16 as effector. The results showed a strong binding affinity of ScbZIP16 protein on promoters of *ScGA20 Oxidase* and *Scent-IKH* harbouring G-box motives in the presence of aureobasidin A (Figure 5B). To further confirm this interaction, a luciferase reporter assay has been performed and results suggested that ScbZIP16 efficiently bound on the promoter of *ScGA20 Oxidase* and *Scent-IKH* as indicated in luciferase activity (Figure 5C). Moreover, to test transcriptional activator or repressor nature of ScbZIP16, a dual luciferase interaction assay has been conducted in *N. benthamiana* leaves. The observation of this assay revealed that ScbZIP16 function as a transcriptional repressor for both genes, *Scent-IKH* and *ScGA20 Oxidase* (Figure 5D). The dual luciferase assay is a convenient and reliable technique to measure promoter activity and transcriptional regulation of reporter gene by normalizing with internal control. Altogether, our results explicitly suggested that ScbZIP16 located in nucleus and strongly binds to promoters of *Scent-IKH* and *ScGA20 Oxidase* to negatively regulate transcription of these genes that determine GA3 level which in turn responsible for cell elongation and varying internodal length in sugarcane varieties.

**Figure 5.**
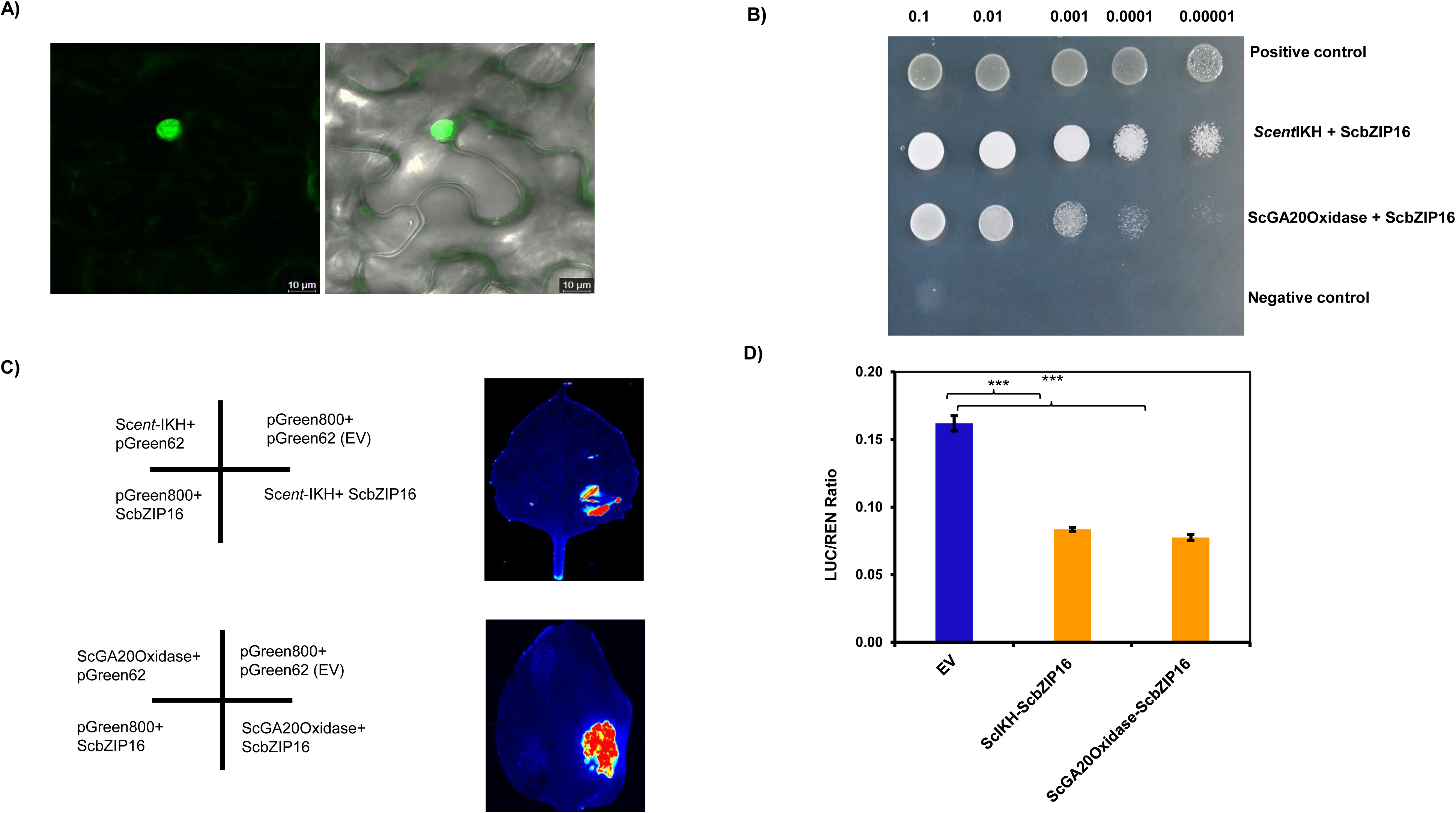
(A) Subcellular localization of ScbZIP16:GFP showing presence in nucleus. (B) Spot assay showing yeast one-hybrid interactions of ScbZIP16 with *proScent-IKH* and *proScGA20 Oxidase* at indicated dilutions. The p53-T7 used as positive control and pLAM-T7 interaction served as negative control. (C) Interaction between bZIP16, *Scent-IKH* and *ScGA20 Oxidase* in luciferase reporter assay. *Agrobacterium* GV3101 strain harbouring indicated combinations was infiltrated into tobacco leaves and luminescence signals were measured 48 h post infiltration. (D) Transcriptional repressor ability of bZIP16 in tobacco leaves. The double-reporter plasmids co-transformed into tobacco leaves by *A. tumefaciens* GV3101, and LUC and REN luciferase activities were assayed post 48 h infiltration. The transcription repression ability of ScbZIP16 was measured by the ratio of LUC to REN. Each value represents the means of three biological replicates, and vertical bars represent the S.D. Significant differences asterisk (***) indicated (*p* < 0.001) by student’s *t*-test, compared with EV. The experiment is repeated three times and data showing here is of one set of experiment.

### 3.6 Expression of *ScbZIP16* altered GA responses in *Arabidopsis*

To prove transcriptional repressor nature, transgenic lines expressing *ScbZIP16* under the control of constitutive promoter have been raised in *Arabidopsis* background. The homozygous lines carrying single copy insertion have been used for the phenotypic variation. Considering GA signal plays an important role in hypocotyl growth and photomorphogenesis, hypocotyl growth assay of transgenic lines and wildtype have been carried out, and quantitative data showed that hypocotyl elongation of dark-grown seedlings of transgenic lines declined significantly relative to wildtype (Figure 6A& B). To know if hypocotyl growth inhibition of transgenics linked to altered GA level, same assay has been conducted either in presence of exogenous GA3 (10 µM) or paclobutrazol (PAC), a potent inhibitor of GA biosynthesis in plant. Exogenous GA3 supplementation increased hypocotyl length of all genotypes which regained to equivalent of the wildtype whereas progressively decreased by increasing dose of PAC in the media (Figure 6A&B). The addition of PAC reduced endogenous GA biosynthesis and abolished hypocotyl length difference between transgenic lines and wildtype suggesting a major role of GA in cell elongation of hypocotyl (Figure 6A&B). The hypocotyl elongation is the outcome of a complex interplay of numerous regulators of photomorphogenesis including GA pathway. Given that AtbZIP16 of *Arabidopsis* interacted with cryptochrome 1 (CRY1), a blue light photoreceptor and involved in blue light (BL) induced photomorphogenesis (Norén et al. 2023). We next investigated effect of BL on hypocotyl growth of transgenic lines and data revealed a significant reduction in hypocotyl length compared to Col-0 implicating a direct role of *ScbZIP16* in elongation (FigureS11B & S11C). Given that GA level affects seed germination and together with other environmental cues such as photoperiod and light quality regulate vegetative growth to reproductive phase transition. We next examined transgenic plant growth from seed germination to attaining maturity. As expected, there was significant decline in seed germination of genotypes at 24 and 36 hrs post sowing relative to wildtype (FigureS11A). Similarly, transgenic lines have displayed a notable delay in flowering time and growth compared to Col-0 suggesting expression of ScbZIP16 altered GA metabolism in Arabidopsis (Figure 6C & FigureS12B).

**Figure 6.**
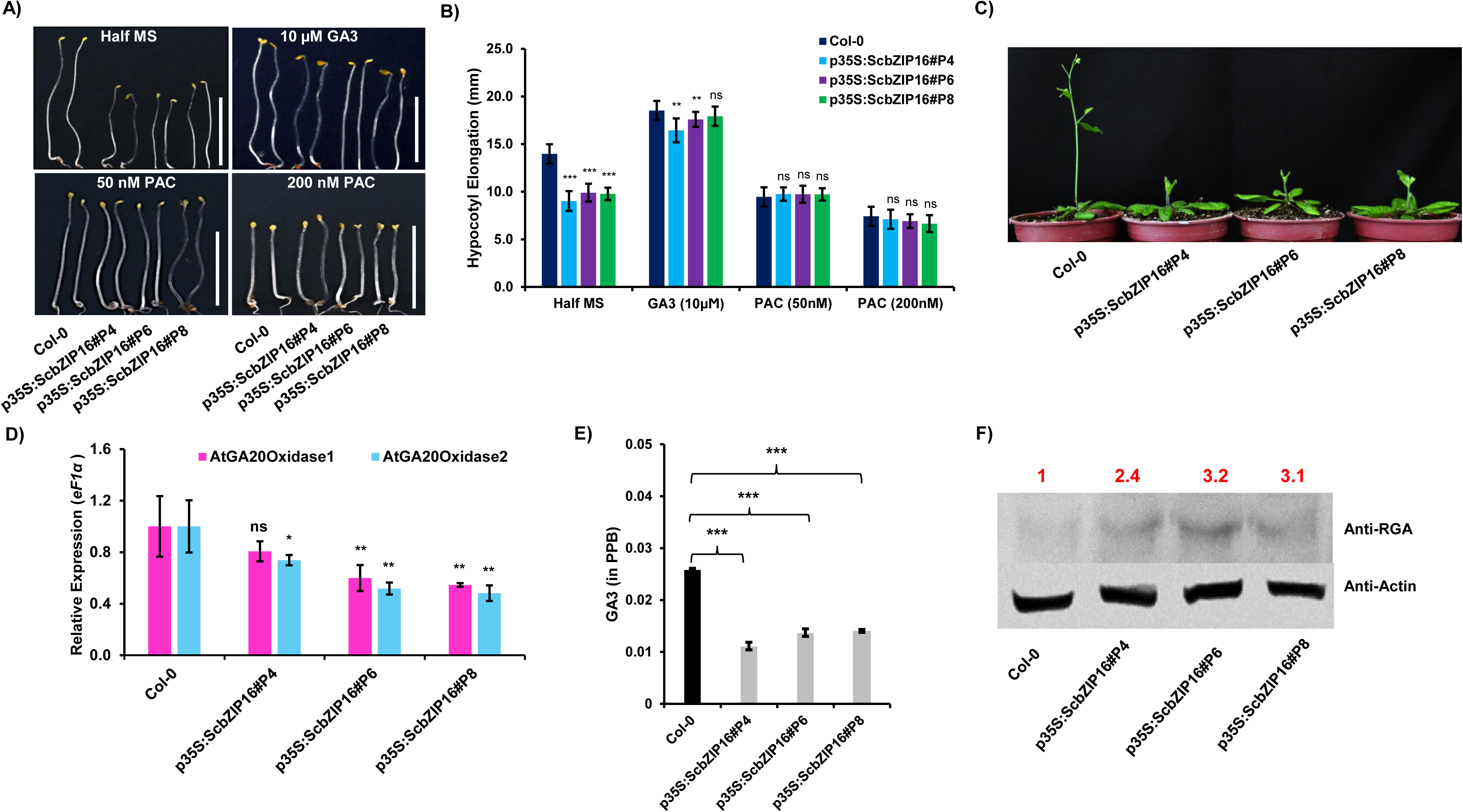
(A) Hypocotyl growth of *ScbZIP16* expressing transgenic lines and Col-0 on half strength MS, supplemented with GA3 and PAC. (B) Hypocotyl elongation of transgenic lines and Col-0 seedling. (C) Flowering delay phenotype of wildtype and transgenic lines (25 days old plant). (D) Quantitative real time PCR analysis of *AtGA20 Oxidase* 1 and *AtGA20 Oxidase* 2 of mature plants (25-day old). (E) GA3 estimation (per 100mg) in transgenic lines and Col-0 of mature plants (25-day old). (F) Western blot assay of RGA in transgenic lines and Col-0 of mature plants (25-day old). Each value represents the means of three biological replicates, and vertical bars represent the S.D. Significant differences asterisk (***) indicates (*p* < 0.001), (**) indicates *p* < 0.001, (*) indicates *p* < 0.05 and ns indicates non-significant. The significant difference calculated using the one-way *ANOVA* (Dunnett test), compared with Col-0.

### 3.7 *ScbZIP16* attenuates GA biosynthesis and signalling in transgenic *Arabidopsis*

In order to know molecular reason, we measured transcripts level of GA biosynthesis genes *AtGA20 Oxidase 1* and *AtGA20 Oxidase 2* in transgenic plants. The qRT-PCR data showed that expression of these genes significantly decreased in the transgenic lines compared to wildtype indicating insufficiency of GA in these lines (Figure 6D). To confirm such observation, GA level has been measured in transgenic plants using LC-MS technique in the rosette of mature plants. Consistent with the previous findings, transgenic lines found to contain significantly less GA3 compared to wild type plants (Figure 6E). DELLA proteins are important regulators of various developmental processes including cell elongation, seed germination and flowering. DELLA proteins activity is determined by GA level in plant. We then decided to measure RGA1 level through immunoblot assay. The data clearly shown that RGA1 protein was stabilized and accumulated at higher amount in transgenic lines corresponding to wildtype (Figure 6F). Altogether, observations of this section established function of *ScbZIP16* as transcriptional repressor of GA biosynthesis and signalling pathway in *Arabidopsis*.

## 4 Discussion

Achieving sugarcane related energy demand, a comprehensive system-based approach integrating sustainable agro-practices and continuous efforts on improving cultivars are absolute necessities. Nearly twelve years timeframe is required to release a new commercial variety after initial crosses covering numerous bottlenecks in conventional sugarcane breeding (Botha and Marquardt, 2024). Advance breeding includes techniques like precision breeding, molecular markers, transgenics, and gene editing for developing new varieties by manipulating key regulatory genes and associated traits. At present, complete knowledge gap exists related to molecular mechanisms and metabolic changes occur in the developing internodes of sugarcane which is not only act as sink of sugar but also linked to stalk biomass. We characterize *ScbZIP16* role as negative regulator of internode elongation by altering endogenous GA pathway in sugarcane and proposing it as a prominent target for sugarcane improvement future strategies.

CoLk16202 found to have longer internode with significantly lesser number of nodes, culm length, cane girth, leaf and stalk biomass compared with CoLk14201 suggesting it possess distinct characteristics (Table 1). Genome size of studied varieties is equivalent to ~10 Gb comparable to early developed Co8371 (Bhima) and CoSe01424 varieties, however, difference of nearly 200 Mb could justify morphological uniqueness (Mondal et al. 2017; D’Ario et al. 2021). Chlorophyll content is significantly higher in CoLk16202 indicating higher photosynthetic rate and efficiency relative to CoLk14201. Chlorophyll fluorescence of plant indicates light energy to chemical energy conversion ability which is crucial for plant growth and productivity. A recent study has shown that *LIMYB* signalling module negatively regulates photosynthesis by repressing expression of photosynthesis related genes (Ferreira et al. 2025) substantiating the downregulation of *LIMYBs* in CoLk16202 (FigureS8). The intercalary meristem lies within growth ring/node of sugarcane wherein meristematic activities are responsible for cane elongation whereas lateral meristem activity in cambium region increases cane girth due to secondary growth for providing strength and more sugar storage in the stalk. Differential expressions of *CDKs*, *cyclins* and *expansins* explain possible molecular reason for contrasting internode length in sugarcane varieties (Choi et al. 2006; Porceddu et al. 2001). In rice, a study reported that β*-expansin* gene is associated with rapid elongation of internodes under deep-water condition (Lee and Kende, 2001). Similarly, number of genes related to sugar metabolism, signalling and *SWEET* transporters are enriched in DEGs of varietal comparisons suggesting existence of strong corelation between genes expressions and sucrose percentage as well as brix in studied varieties (Table 2; FigureS2& S3).

More expression of genes involved in MEP pathway especially *Scent-IKH* and *ScG20 oxidase* resulting enhanced GA3 accumulation in long internode variety. This observation aligned to a recent study in which overexpression of *ScGA20 Oxidase* causes elevated GA level and accelerated growth of sugarcane (Wang et al. 2025). The similar observations was also recorded in plants such as rice (Qin et al. 2012) and poplar (Cho et al. 2019). The GA3 leads to degradation of DELLA protein, a negative regulator of GA signalling, and overexpression of *DELLA SLR1*-like gene in various plant species caused a delay in growth (Itoh et al. 2005; Li et al. 2015). In this study, we also noticed canonical downregulation of *DELLA SLR1*-like gene in the internode corroborating enhanced GA signalling in long internode variety. The fact that allelic variation/mutation in DELLA protein and plant height have been extensively studied in cereals including wheat during green revolution already established a molecular link between amount of GAs and plant growth (Peng et al. 1999; Sokolowska et al. 2026).

In addition, role of *bZIPs* has been elucidated in shaping plant growth and architecture *via* controlling GA level in tobacco (Fukazawa et al. 2000). Our finding suggest that ScbZIP16 participates in internode elongation through controlling GA pathway indicating that it is a key component of elongation process. Emerging recent evidences suggest that *bZIPs* especially *Hy5*, directly or through interacting with other factors alters gene expressions and ultimately changing endogenous GA level in the tomato (Dong et al. 2023), citrus (Fu et al. 2024) and poplar (Gao et al. 2024) affecting bud dormancy and overall plant growth. An interesting expression pattern of *ScbZIP16* between short and long internode cultivars has been seen in this study led us to postulate that ScbZIP16 could alter GA synthesis in sugarcane. The *in-vivo* interaction experiments in yeast and *N. benthamiana* establish a positive interaction of the ScbZIP16 with *Scent-IKH* and *ScGA20 Oxidase* promoters (Figure 5). In addition, dual luciferase-based transactivation result provides clear evidence that ScbZIP16 act as a transcriptional repressor for *Scent*-*IKH* and *ScGA20 Oxidase* which directly responsible for varying GA3 levels in sugarcane varieties. The heterologous expression retains same GA biosynthesis repressor nature of *ScbZIP16* in *Arabidopsis* as well. The hypocotyl growth of transgenic lines is notably reduced either in dark or BL conditions compared to wildtype. The observations of hypocotyl growth assay in the presence of GA3 and PAC clearly confirms that more GA signal promoted hypocotyl growth and less GAs cause negative growth in *Arabidopsis* (Alabadí et al. 2004) (Steber and McCourt, 2001; Hsieh & Hsieh, 2012). Similar to sugarcane, four core bZIPs binding motives has been identified in promoters *AtGA20 oxidase 1* and *AtGA20 Oxidase 2* (FigureS12A). The GA regulates not only seed germination but also floral development in DELLA dependent manner in coordination with other environmental signals. In general, DELLA proteins are known for repressors of GA signalling and GAs stimulates DELLA degradation to promote growth related activities in plants (Silverstone et al. 2001; Davière et al. 2016). Among five *DELLAs* in the *Arabidopsis* genome, RGA and GAI control cell expansion and cell division in hypocotyl whereas RGA together with RGL1 and RGL2 induce vegetative to floral transition phase (Dill and Sun, 2001; Daviere et al. 2014; Tyler et al. 2004). Reduced GA3 level in transgenic lines, stabilization of DELLA protein and delay in flowering time provides multiple line of evidences confirming that *ScbZIP16* suppresses GA pathway in *Arabidopsis* and sugarcane (Figure 6).

In summary, our study revealed a mechanism whereby a ScbZIP16 regulated internode growth and plant development by affecting the biosynthesis of GAs and DELLA accumulation in plants (Figure 7). This regulatory module might benefit for improving stalk growth as low sugarcane yield per hectare is a global challenge compared to theoretical yield potential of sugarcane. This study of *ScbZIP16* not only provides a candidate for sugarcane breeding program but also a new target gene for biotechnological interventions of sugarcane improvement.

**Figure 7.**
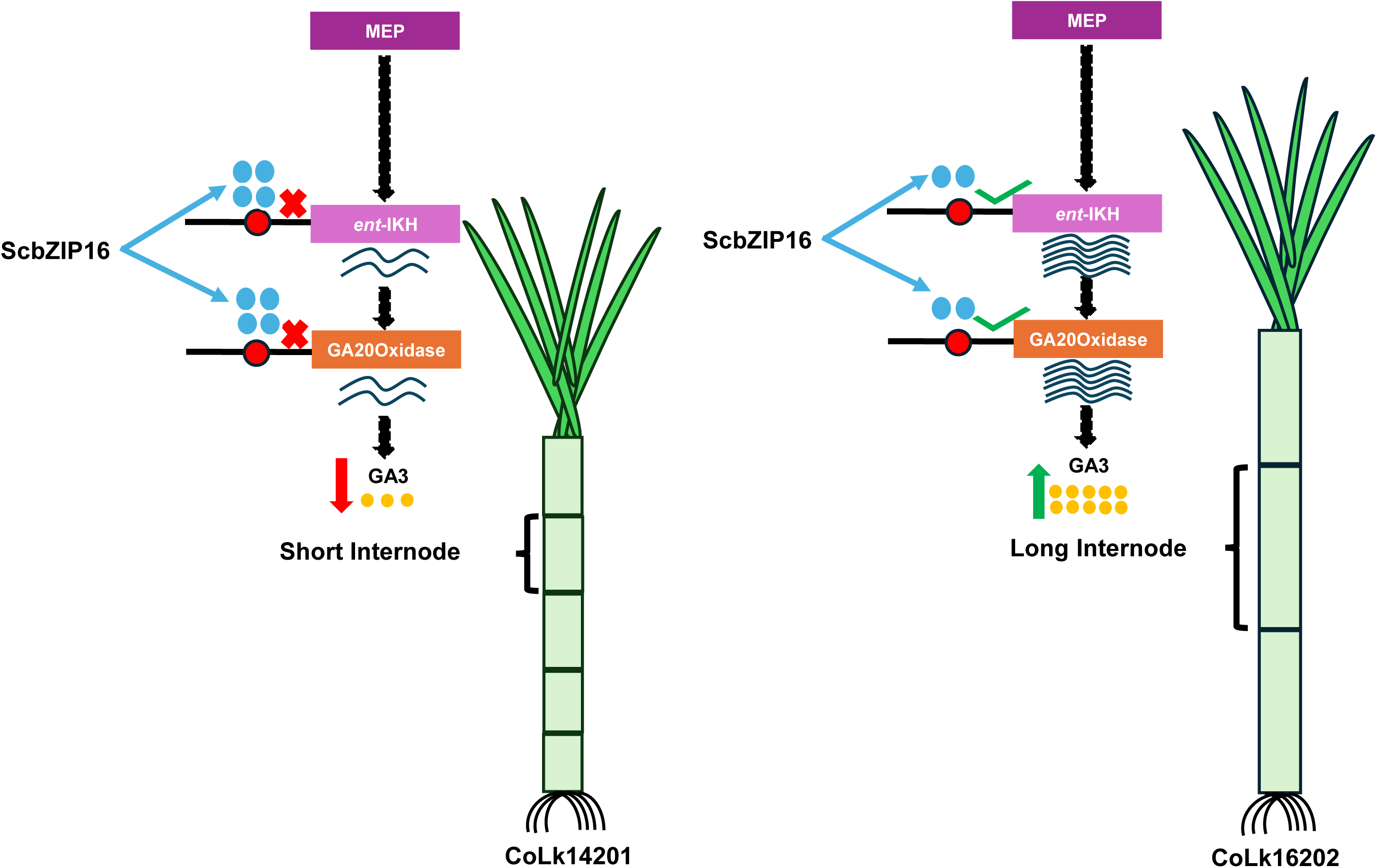
Schematic model showing the regulatory interaction of ScbZIP16 transcription factor with *proScent-IKH* and *proScGA20 Oxidase* highlighting molecular module of internodal elongation in short and long internode sugarcane varieties.

## Acknowledgements

SS thanks to ANRF for providing National Post-Doctoral Fellowship. SS and SJ thank to Council of Scientific and Industrial research and University Grants Commission, India, respectively for senior research fellowships. Authors are also thankful to Dr. Prashant Misra, CSIR-Indian Institute of Integrative Medicine, Jammu for sharing the vectors used in luciferase assays; Dr. Jaroslav Doležel, Laboratory of Molecular Cytogenetics and Cytometry, Institute of Experimental Botany, Olomouc, Czech Republic for the gifting seeds of *P. sativum* L Citrad, and Dr. Satya Narayan Jena, CSIR-NBRI, Lucknow for his help in flow cytometry. This work was partially supported by the National Laboratories Scheme under the ULIP scheme ‘Conservation and utilization of botanical wealth of India (NBRI-BOT) [Grant Number MLP002628].

## Authors’ contributions

SS carried out the experiments, recorded the observations, performed the bioinformatics analysis and drafted the manuscript. SS and RG performed the experiments and recorded the observations. VT and MHA analysed the transcriptome data. SJ helped in bioinformatics analysis and metabolite analysis. MKS performed the physiological and biochemical experiments. AKS and MT conceptualized the study, supervised experiments, drafted and edited the manuscript.

## Data Availability Statement

The transcriptome data of the present study has been submitted to the NCBI SRA database under the BioProject accession number PRJNA1429795.

## Supporting Information

The data that supports the findings of this study are available in the supporting information files of this article.

