## Supporting Information Figures for "Sugarcane *bZIP16* regulates internode elongation by targeting gibberellic acid biosynthesis pathway"

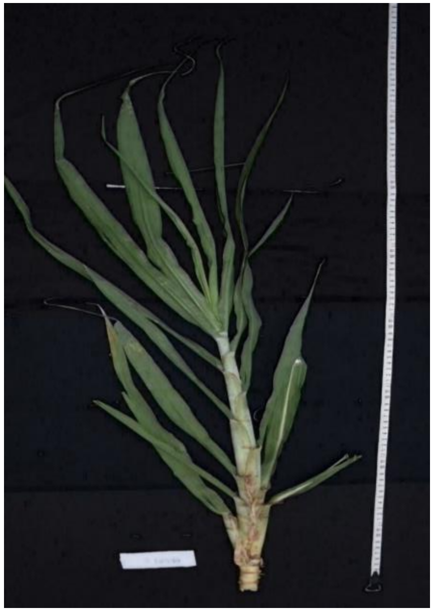

CoLk14201

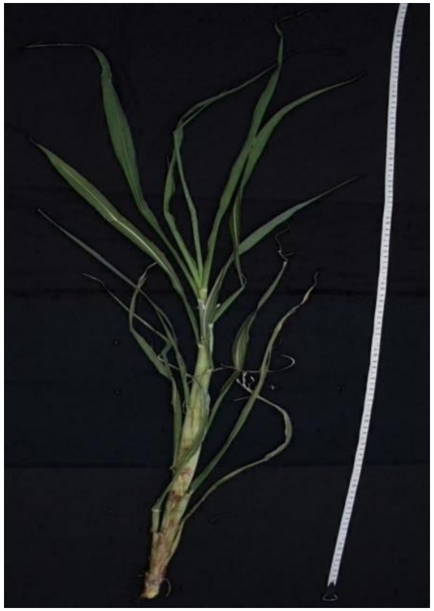

CoLk16202

**Figure S1** Image of whole plant of sugarcane varieties at early stage (S1) when node formation initiated.

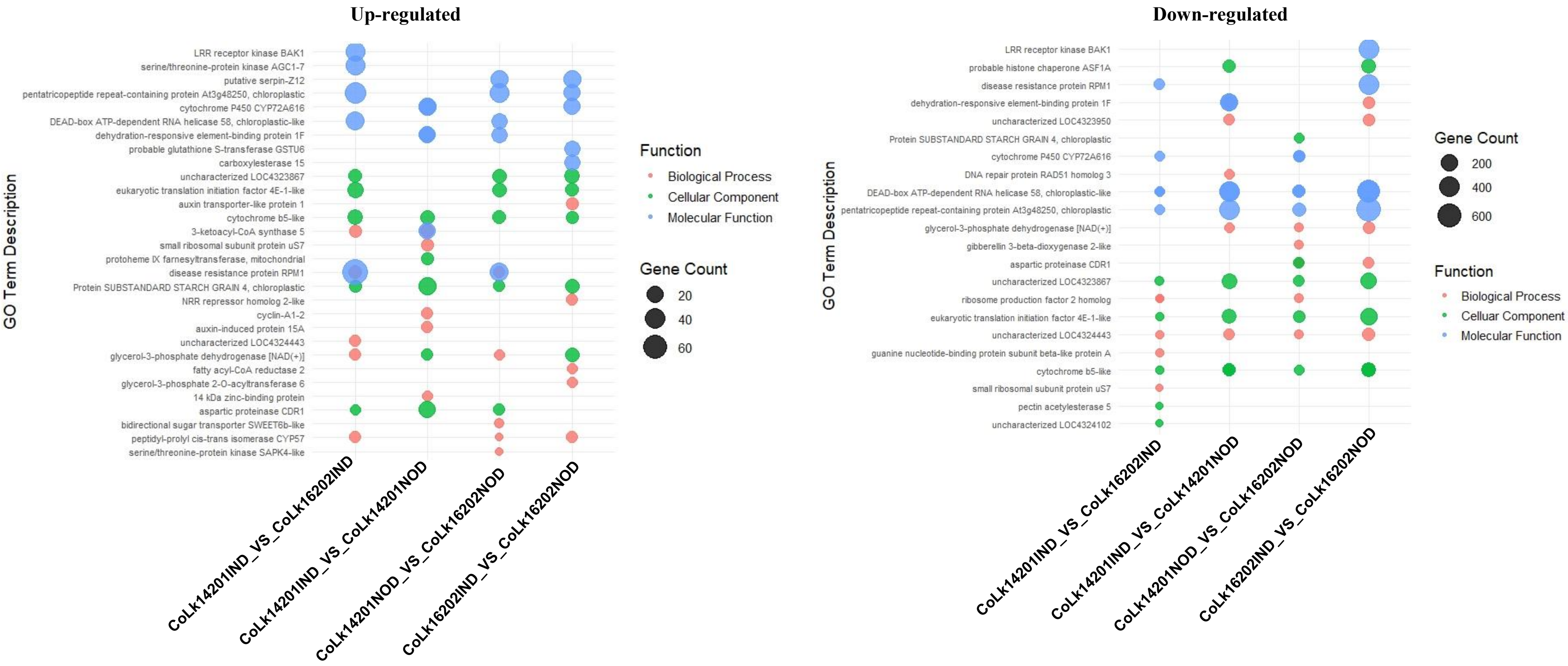


**Figure S2** GO function bubble plot of DEGs across different comparisons in sugarcane varieties at early growth stage (S1) when node formation initiated.

**Up-regulated**

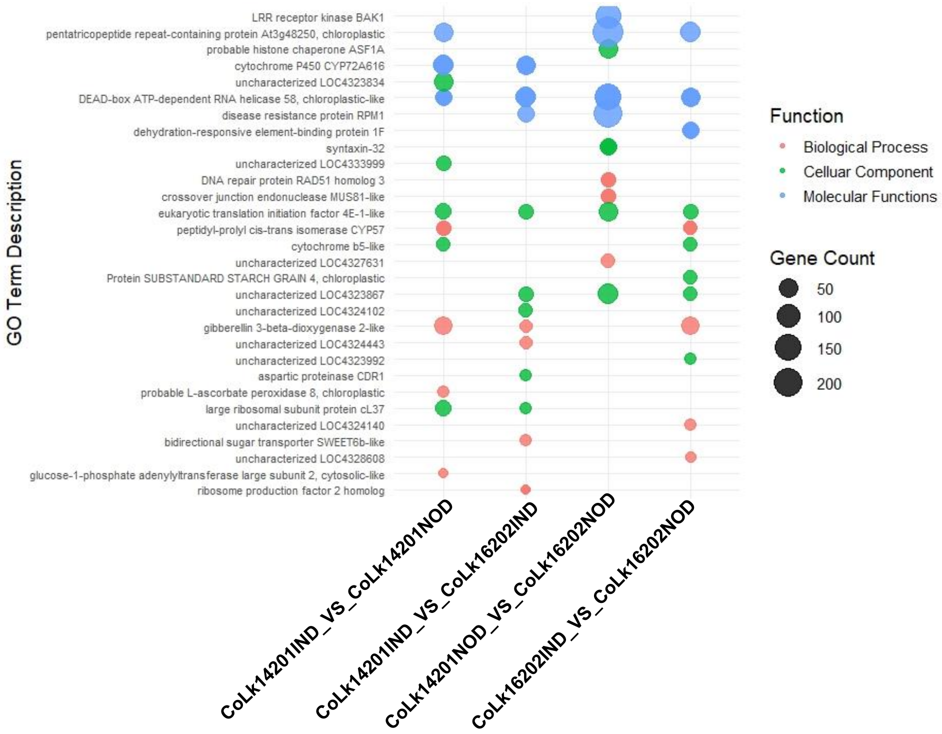


**Down-regulated**

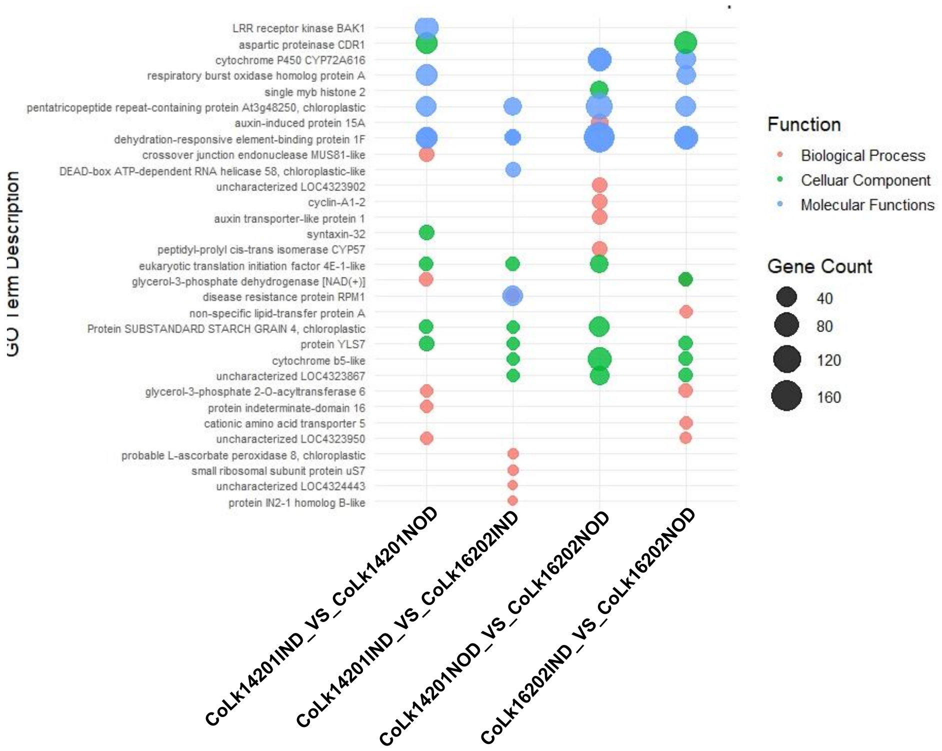


**Figure S3** GO function bubble plot of DEGs across different comparisons in sugarcane varieties at mature growth stage (S2) when cane growth terminated.

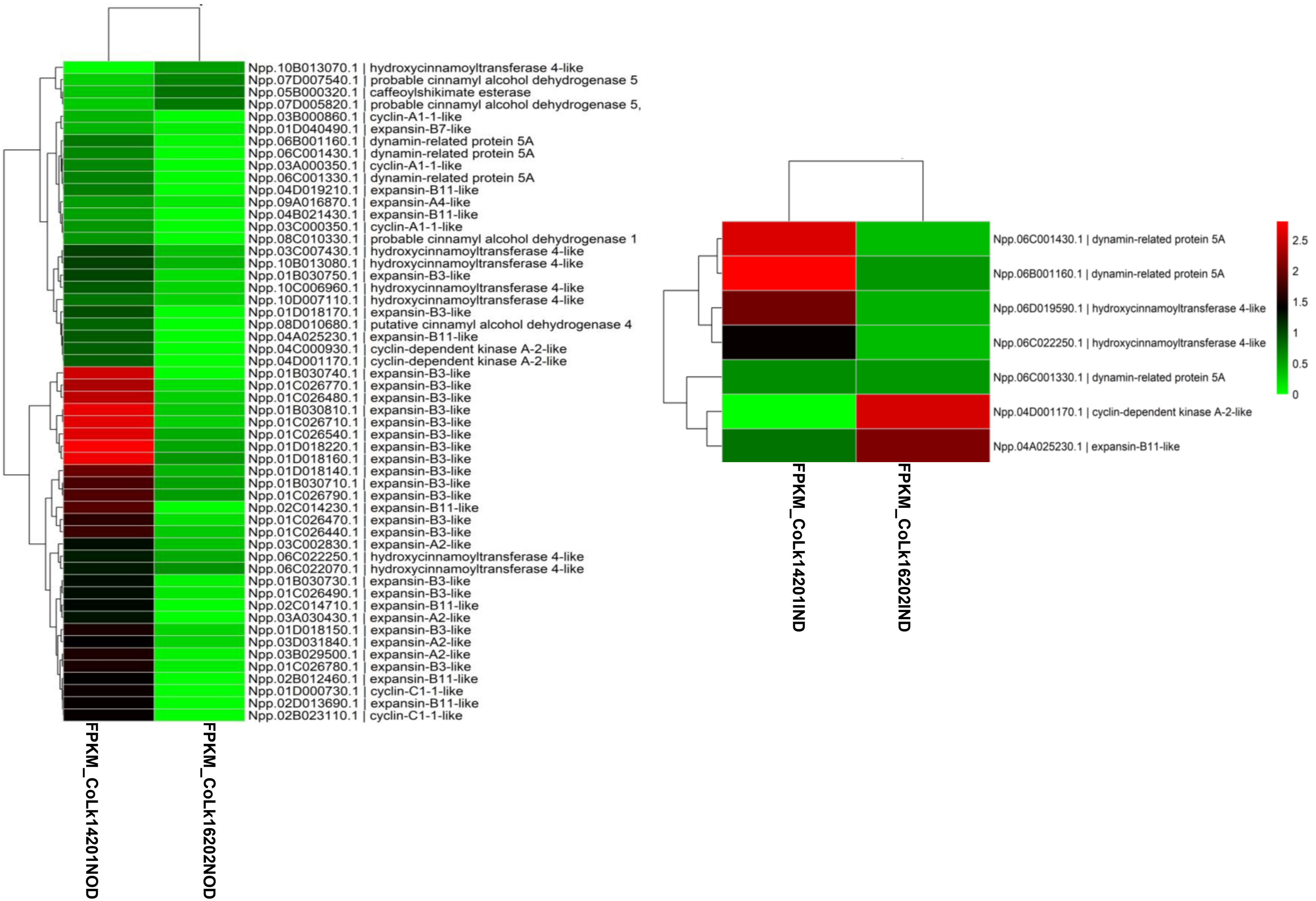


**Figure S4** Expression of growth and cell division related genes in sugarcane varieties at early stage (S1) when node formation initiated.

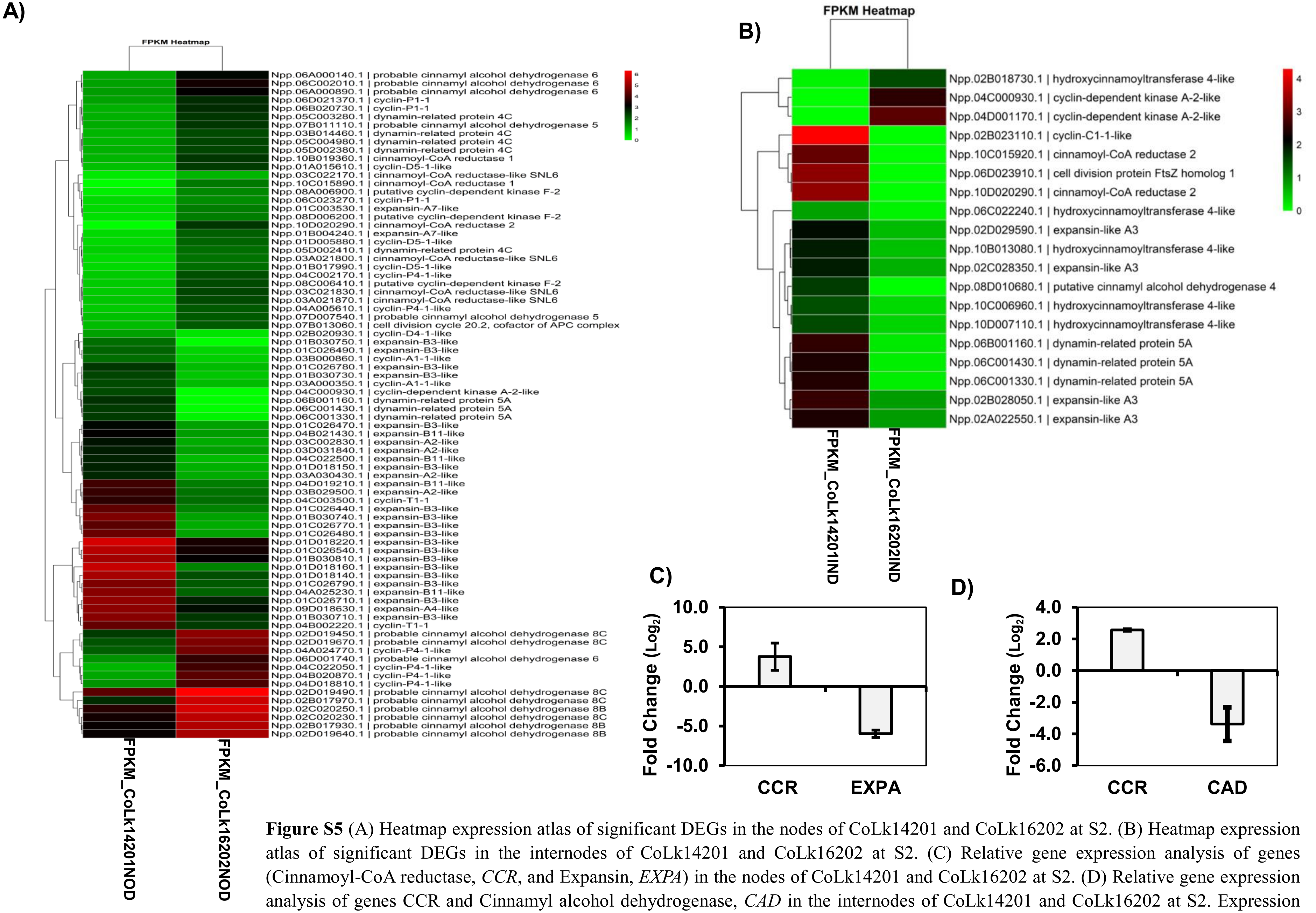


**Figure S5** (A) Heatmap expression atlas of significant DEGs in the nodes of CoLk14201 and CoLk16202 at S2. (B) Heatmap expression atlas of significant DEGs in the internodes of CoLk14201 and CoLk16202 at S2. (C) Relative gene expression analysis of genes (Cinnamoyl-CoA reductase, *CCR*, and Expansin, *EXPA*) in the nodes of CoLk14201 and CoLk16202 at S2. (D) Relative gene expression analysis of genes CCR and Cinnamyl alcohol dehydrogenase, *CAD* in the internodes of CoLk14201 and CoLk16202 at S2. Expression analysis was performed using pooled samples of three independent biological and three technical replicates. Vertical bars represent the S.D

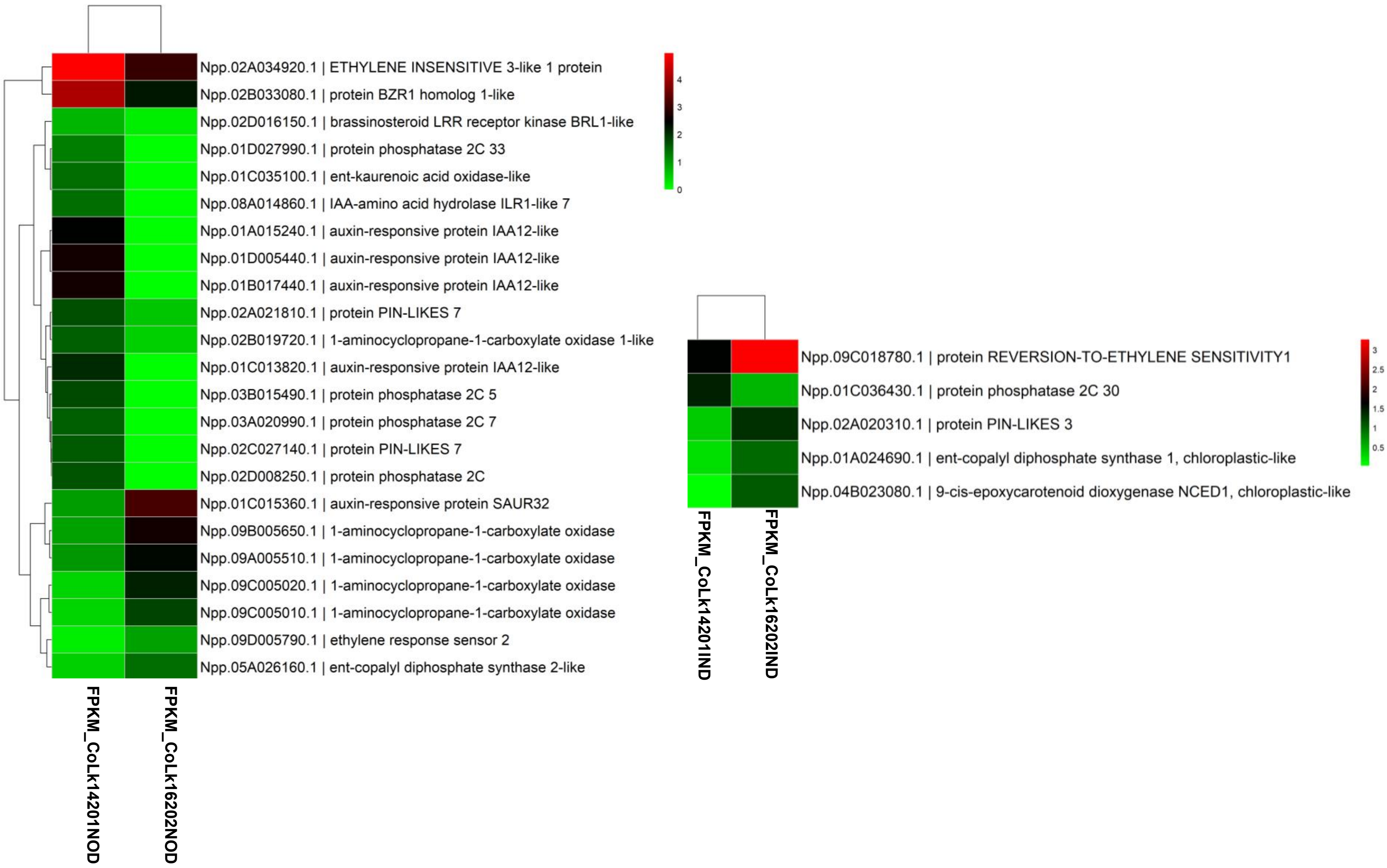


**Figure S6** Heatmap expression of phytohormones related genes enriched in DEGs of node and internode at early growth stage (S1).

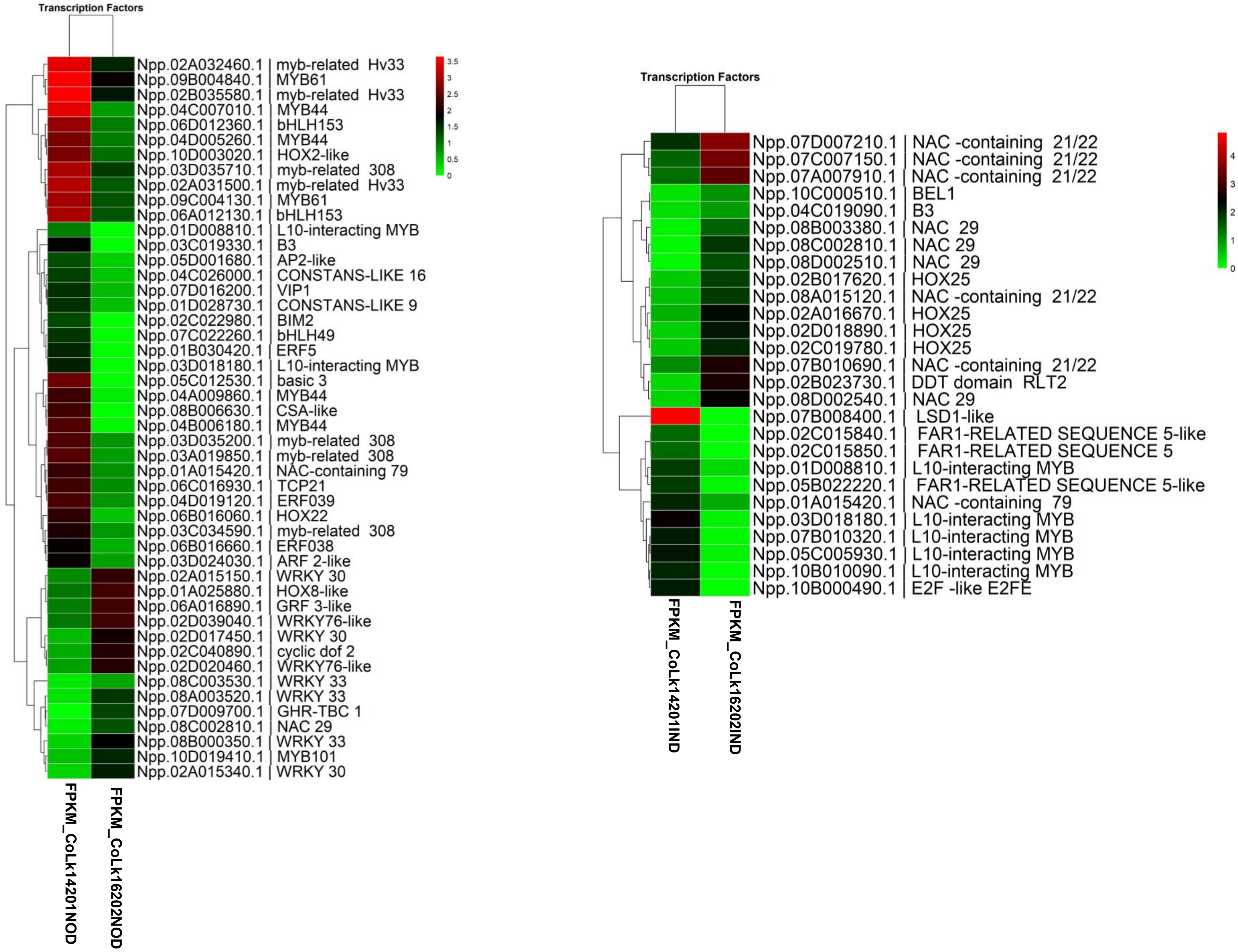


**Figure S7** TFs related genes in sugarcane varieties at early stage (S1) when node formation initiated.

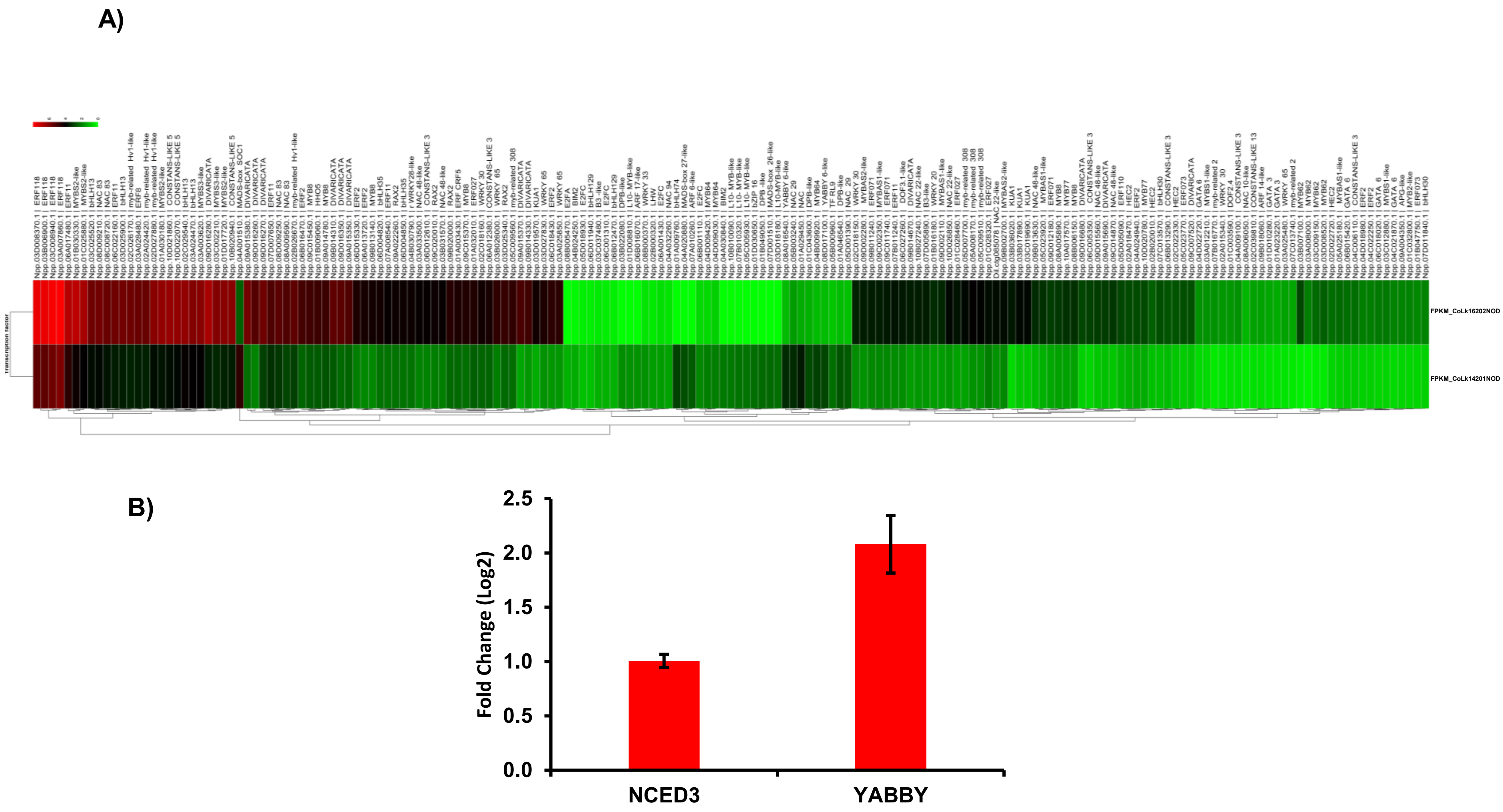


**Figure S8 A)** Heatmap expression of transcription factor related genes enriched in DEGs at mature growth stage (S2) in node.

**B)** Relative gene expression analysis of 9-cis-epoxycarotenoid dioxygenase, *NCED3 and YABBY* in the node of CoLk14201 vs CoLk16202 which was analysed using pooled samples of three independent biological and three technical replicates

A)

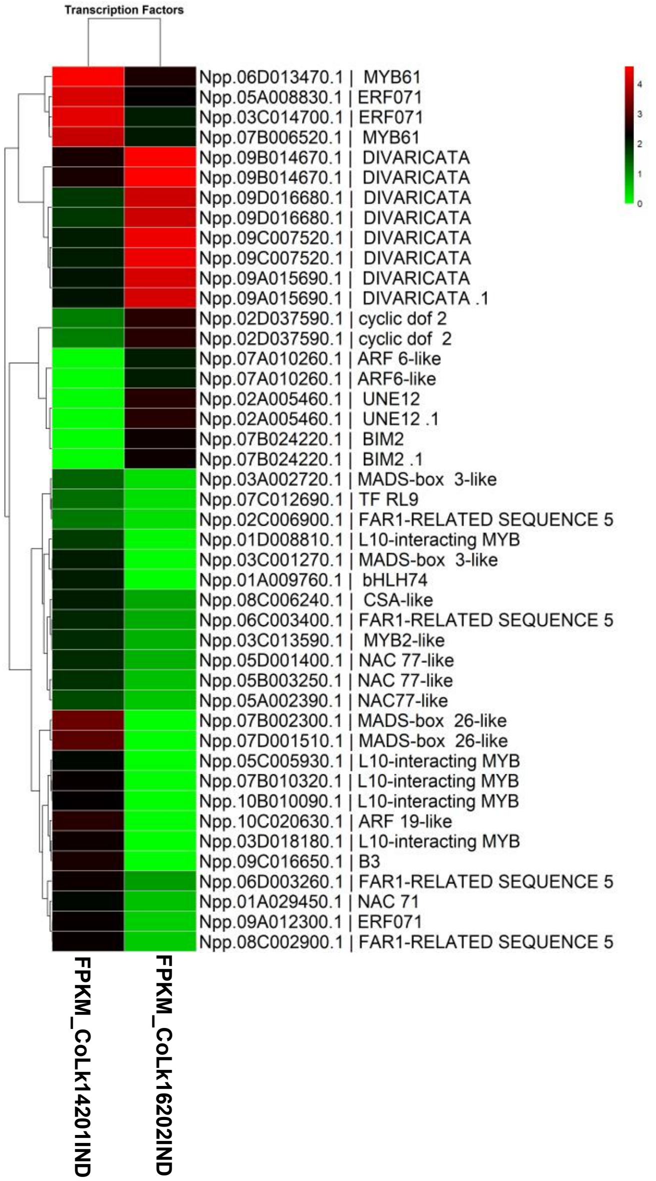


B)

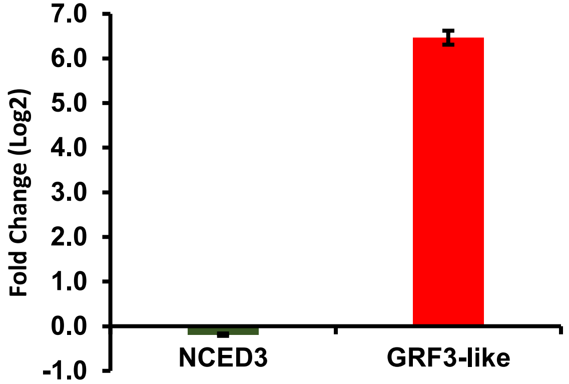


**Figure S9 (A)** Heatmap expression of transcription factor related genes enriched in DEGs in internode at mature stage (S2).

**(B)** Relative gene expression analysis of 9-cis-epoxycarotenoid dioxygenase, *NCED3* and *growth regulatory factor 3-like, GRF3-like* in the internode of CoLk14201 vs CoLk16202 which was analysed using pooled samples of three independent biological and three technical replicates

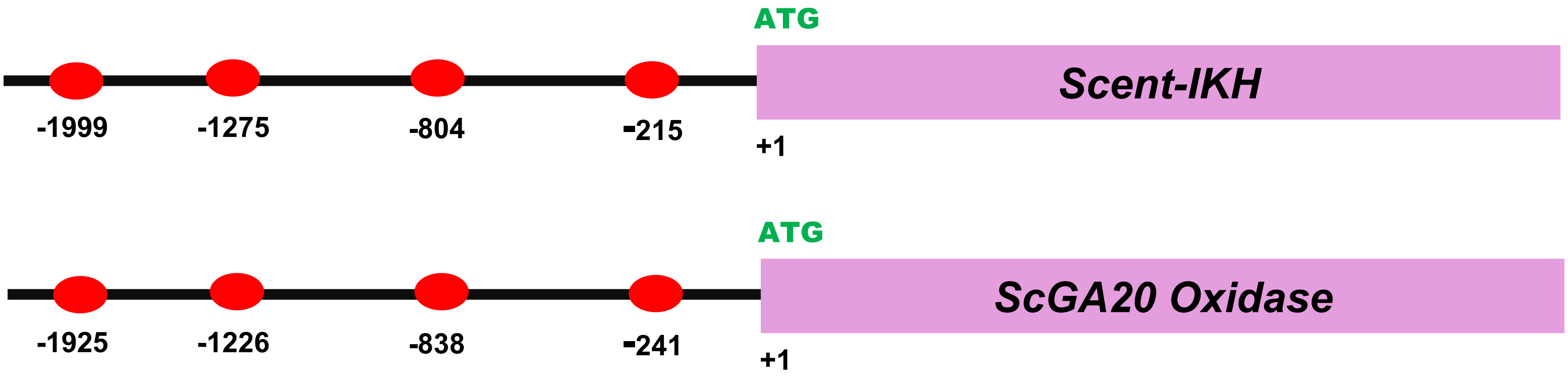


**Figure S10** Representation of the position of bZIP binding motifs in promoter sequences of *Scent-IKH* and *Sc20 Oxidase*.

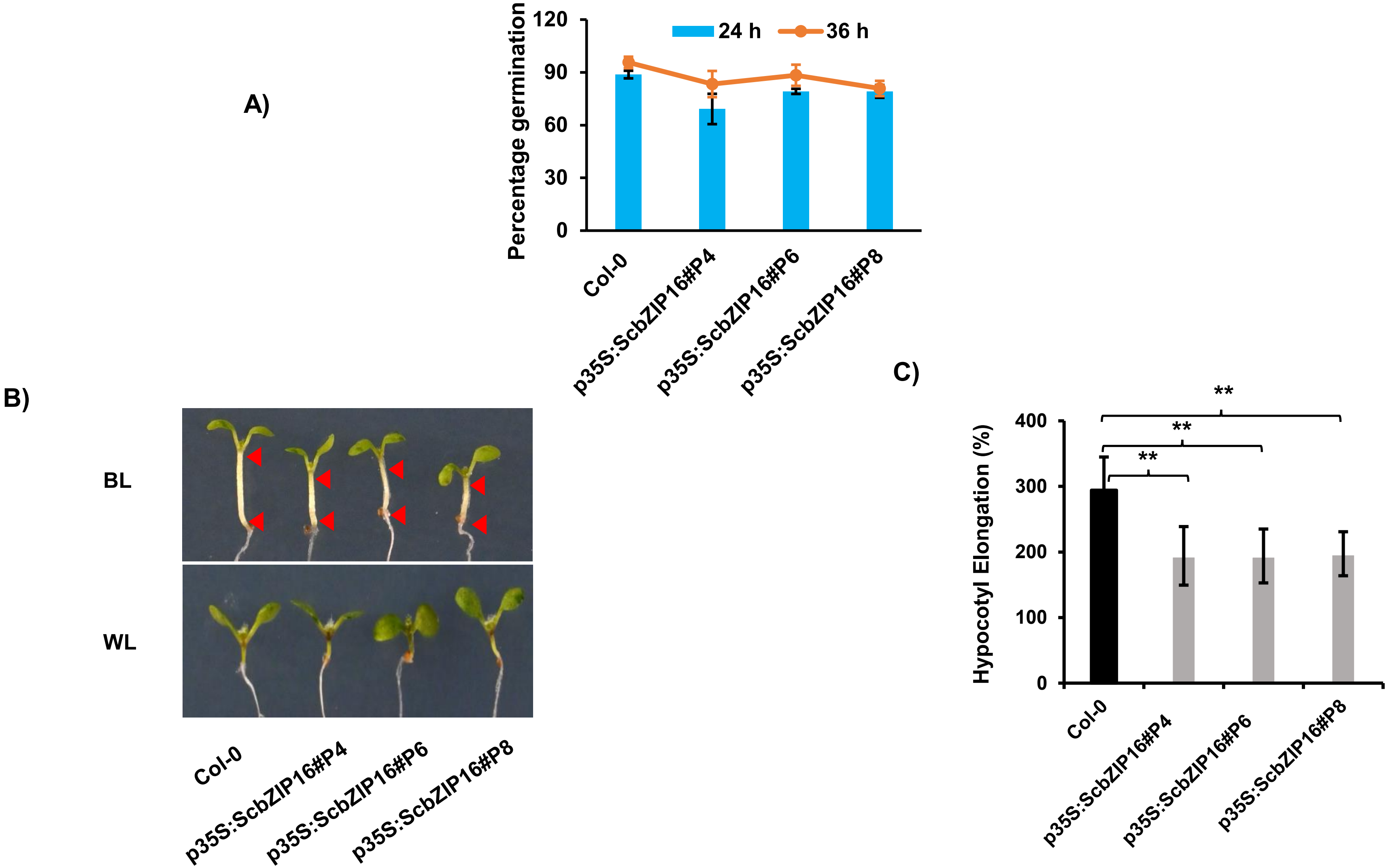


**Figure S11 (A)** Germination percentage of transgenic lines expressing *ScbZIP16* and Col-0 growing under white light condition (114µM/m$^2$/s) at indicated time interval (B) Hypocotyl elongation of transgenic lines and Col-0 under blue light (35µM/m$^2$/s$^{-1}$) for five days. The plants were initially allowed to grow under white light condition for two days, and then shifted to blue light regime (C) The measurement of hypocotyl elongation percentage of transgenic lines and Col-0 under Blue light. Each value represents the means of three biological replicates, and vertical bars represent the S.D. **Significant differences in values (**$p < 0.01$) by one-way Anova (Dunnett test), compared with Col-0. The experiment is repeated three times and data showing here is of one set of experiment.

A)

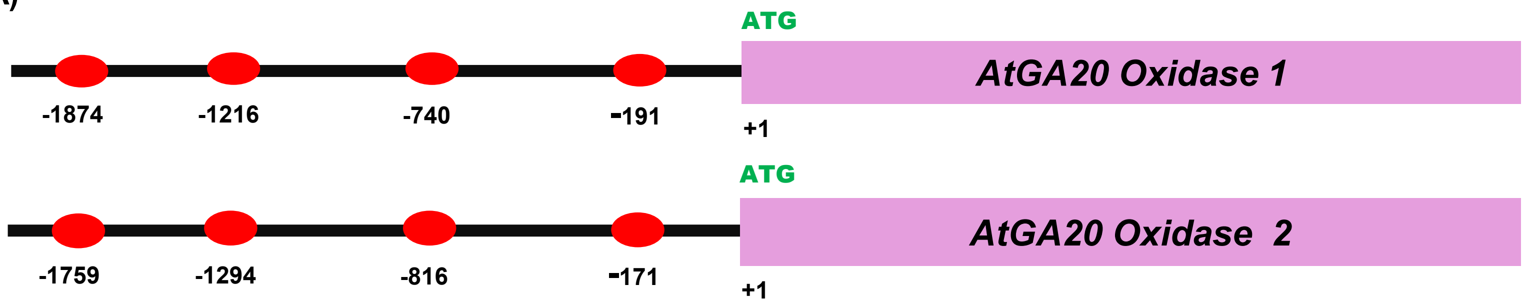


B)

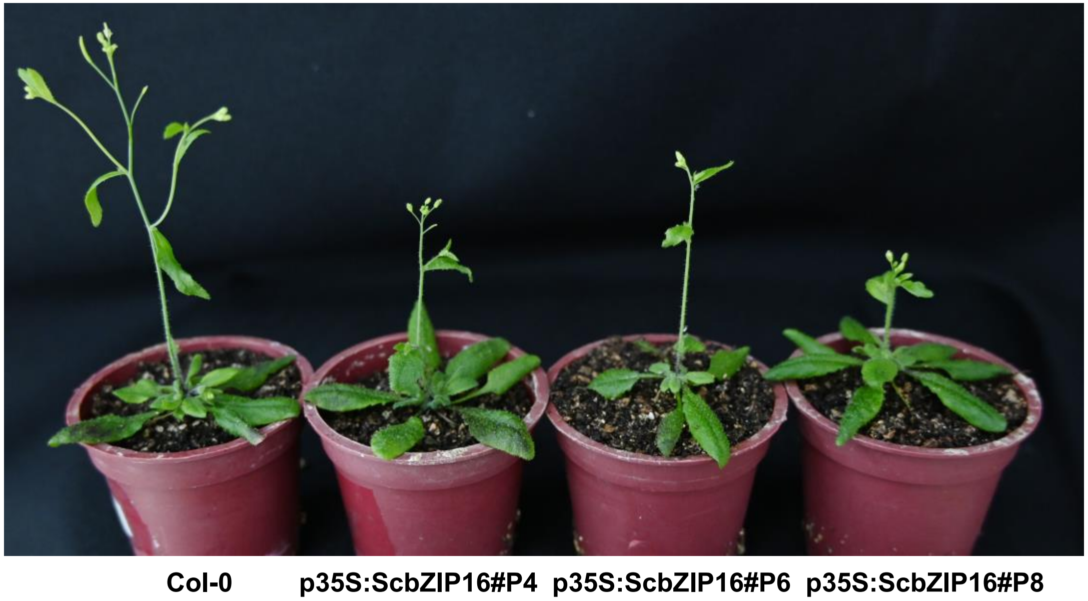


**Figure S12 (A)** Representation of the position of bZIP binding motifs in promoter sequences of *AtGA20 Oxidase 1* and, *AtGA20 Oxidase 2*. (**B)** The image of mature plant (30 days old) of Col-0 and transgenic lines showing delayed flowering time under standard growth conditions in long day photoperiod. The experiment is repeated three times and image showing here is of one set of experiment.
